# The TR locus annotation and characteristics of Rhinolophus ferrumequinum

**DOI:** 10.1101/2021.06.15.448458

**Authors:** Hao Zhou, Long Ma, Longyu Liu, Xinsheng Yao

## Abstract

T cell antigen receptors (TCRs) in vertebrate could be divided into αβ or γδ, which are encoded by TRA/D, TRG and TRB loci respectively. TCRs play a central role in mammal cellular immunity, which are functionally produced by rearrangement of V gene, D gene, J gene and C gene in the loci. Bat is the only mammal with flying ability, and is considered as the main host of zoonotic virus, which occupies an important position in the field of public health. At present, little is known about the composition of bat TRs gene. According to the whole genome sequencing results of the Rhinolophus ferrumequinum, and referring to the TR/IG annotation rules formulated by IMGT. We make a complete annotation on the TRA/D, TRG and TRB loci of the Rhinolophus ferrumequinum. A total of 128 V segments, 3 D segments, 85 J segments and 6 C segments were annotated, in addition to compared with the known mammalian, the characteristics of the TRs locus and germline genes of the Rhinolophus ferrumequinum were analyzed.

## 1 Introduction

Vertebrate T cells participate in adaptive immune response via their diversified TCRs on the surface. TCRs composed of two polymorphic chains, which can be divided into two types: αβ polypeptide chain and γδ polypeptide chain, which are encoded by four loci TRA/D, TRB and TRG, respectively. Each polypeptide chain contains a variable region and a constant region, which are rearranged by germline V (variable, V) gene and/or D (Diversity, D) gene and J(Joining, J) gene[1, 2]. The recombination process of TCR is conducted by the product of RAG (Recombination activating gene, RAG) 1/2 gene, by making double-strand breaks at the recombination signal sequence, and then shearing and insert to rearrange the V(D)J gene[3]. IG and TR loci contain hundreds of germline genes, humans contain 608-665 BCR and TCR genes, and mice contain more than 800 BCR and TCR genes, which can be divided into functional genes, ORFs and pseudogenes according to functionality[4]. The annotation of TR and IG loci is challenging, because germline genes do not have the classical intron/exon structure that can be detected by standard annotation software, and most of the sequences are very short, such as J gene between 40-60bp, especially D gene only about 10bp. IMGT database (IMGT Aide-mémoire, http://www.imgt.org/) is the authoritative database of immunogenetics and immunoinformatics, which describes the accurate annotation rules for TR and IG genes in detail. In the process of locus annotation of TR or IG: firstly, the identification of V, D, J, and C genes is the most important part; then a complete description of each gene, for example, for complete V gene segment includes: L-part (L-part1+L-part2)+V-region+V-RSS; next is the functional determination of germline genes, which can be divided into 3 types according to IMGT:(1) Functionality gene (Functionality, F) means that there is an open reading frame and there is no defect of stop codon and splice site; according to the severity of the identified defect, it can be defined as (2) ORF (Open reading frame, ORF) and (3) P (Pseudogene, P); the final step is to classify the group or subgroup according to the nucleotide similarity and the location of cluster [5-7].

Vertebrate contain 4 TR loci. At present, TR loci have been completely or partially annotated in humans, monkeys, mouse, cattle and other species (http://www.imgt.org/IMGTrepertoire/LocusGenes/#C), the structural characteristics of different loci are also summarized. For species after teleost, TRA and TRD locus are located in the same loci. TRA locus and TRD locus share part of V gene, but independent J and C genes. The mammalian TRD locus is usually located downstream of the TRAV gene, upstream of the TRAJ gene, there is always a TRDV gene with the opposite transcription direction downstream of the TRDC gene[8]. TRG locus usually has two structures: one is the typical form of V-J-C translocon, where the replication occurs in J-C clusters, such as humans, rabbits, possums, etc.; the other is that V-J-C clusters are formed in multiple due to genome duplication. Structures, such as cattle, dog, mouse, etc., are quite different in the V-J-C cluster and J-C cluster of the TRG locus among different species [9]. The structure of the TRB locus in mammals is relatively conservative. A common feature is semi-cluster. The upstream usually contains multiple V genes with varying numbers, and there are several D-J-C clusters distributed downstream. The number varies from species to species. The D-J-C cluster increases the number of germline genes pool that could be used for recombination [10].

Although there are many kinds of bats (more than 5,000), there is few changes in the process of evolution. Bats account for 20% of all mammals and have settled in six continents[11-13]. Bats are the host of variety viruses and carry a large number of virulent viruses. There have been many studies indicate that bats will not cause clinical symptoms for these viruses [14-16]. At present, although the reduction and increase of a large number of immune-related genes have been identified in a few bats, due to the lack of experimental models or related reagents, the research on adaptive immunity of bats is quite limited [17]. Major mammalian antibody subclasses have been detected in bats, including IgA, IgE, IgM and IgG [18, 19]. Early studies reported that the scale level and duration of antibody reaction of bats to antigens may be lower than those seen in conventional experimental animals [20, 21], but the specific role of antibodies in bats in virus infection is still unclear. At the genome level, bats seem to have more BCR germline genes than humans, which may provide more antigen specificity for the naive B-cell receptor. Meanwhile, no somatic high-frequency mutation is found in the small brown bat, which also indicates that bats may rely more on germline pools to respond infection [22, 23].

At present, the genomes and transcriptomes of at least 18 species of bats could be obtained in the database [24, 25], including Pteropodidae: Pteropus vampyrus (GCA_000151845.2), Pteropus alecto (GCA_000325575.1), and Vespertilionidae: Myotis lucifugus (GCA_000147115.1), Myotis davidii (GCA_000327345.1) and Rhinolophidae: Rhinolophus ferrumequinum (GCA_004115265.3), Rhinolophus sinicus (GCA_001888835.1), etc. However, due to the incompleteness of sequencing and annotation of these bats’ genomes and transcriptome, the systematic and complete annotation of bats’ whole genome has not been completed at present, especially the annotation of bats’ immune genes such as TCR, BCR and MHC is currently blank. According to the annotation rules of IMGT-ONTOLOGY, we annotated TRA/D, TRB, TRG of Rhinolophus ferrumequinum completely, and compared the annotated genes with those of human, mouse, pig, dog, cattle and other species in the IMGT database.

## 2 Methods and Materials

### 2.1 Annotation of the TR locus of Rhinolophus ferrumequinum

According to the whole genome sequence (GCA_004115265.3) shared in NCBI, which was submitted by vertebrate genomes project through whole genome shotgun sequencing, the genome has 28 chromosomes, with a total length of 2075.77Mb, and we could complete the current analysis due to the high quality and coverage of this genome.

We completed the annotation work according to the process of **Figure 1**. The first step is to locate the TR locus in chromosome. According to the borne gene information of mammalian TR loci recorded in IMGT database, TRA/D loci are usually located in OR10G2 (Olfactory receptor family 10 subfamily G member 2, OR10G2) gene and DAD1(Defender against cell death 1, DAD1). The TRG locus is located between AMPH (Amphiphysin, AMPH) gene and STARD3NL (Related to steroidogenic acute regulatory protein D3-n-terminal like, STRAD3NL) gene. TRB locus is located between MOXD2 (Monooxygenase-beta-hydroxylase-like 2, MOXD2) gene and EPHB6 (Ephrin type-b receptor 6, EPHB6) gene.

**Figure.1.**
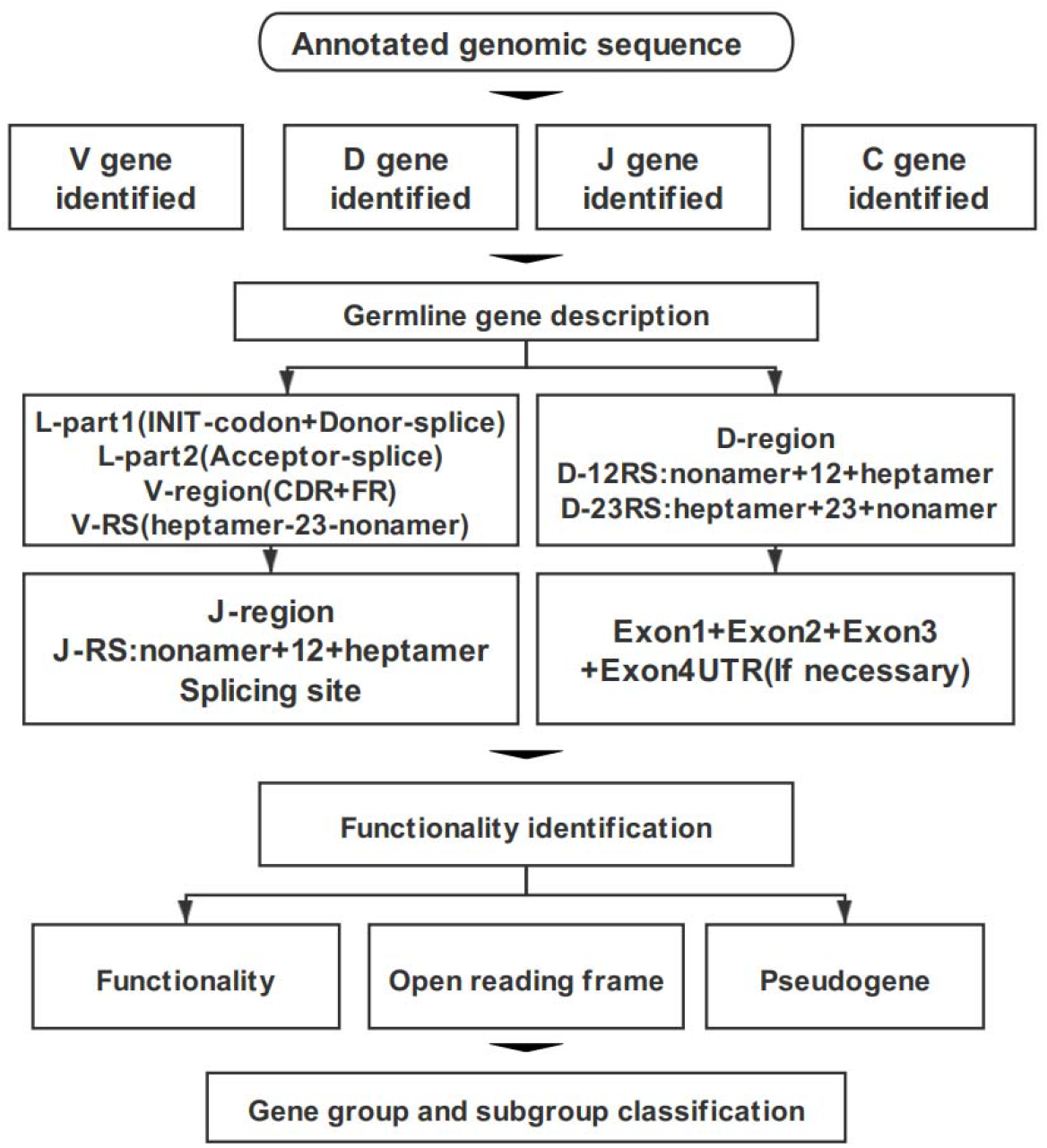
The annotation process of TR locus includes four parts: identification, description, functional determination and family clustering of germline genes.

The second step is the identification of germline genes. We analyzed sequences among the borne genes: we analyzed the V gene based on human and mouse pattern by IgBLAST tool (for the V gene of TRA/D locus, if the output result is TRAV/TRDV, we will annotate it as the V gene shared by TRA and TRD loci). Furthermore, the sequence was imported into Geneious Prime software, and the homology of the sequence was compared by using the germline gene information recorded in IMGT database, including the existing V, D, J and C genes of human, mouse, pig and dog.

The third step is description of germline genes. Here, four parts are described according to the types of germline gene. (1) V gene is composed of 3 parts: L-Part+V-exon+V-RS, L-part contains two segments: L-part1 and L-part2, there is a part of intron between part1 and part2, L-Part1 contains 2 conservative components: Init-codon and Donor-splice, L-part2 contains 1 conservative component: Acceptor-splice, part2 is next to the V-exon part, and V-exon is divided into 6 segments: FR1+CDR1+ FR2+CDR2+FR3+CDR3, and the 3 FR regions contain 3 conservative amino acid positions: 1st-Cys, Conserved-Trp, 2nd-Cys. The last original is 23RSS: V-Heptamer+V-Spacers(23bp)+V-Nonamer. (2) D gene consists of 3 parts: 5’D-RS+D-region+3’D-RS, 5’D-RS includes: 5’D-Nonamer+5’D-Spacers(12bp)+5’ Heptamer, D-region usually consists of about 10-15 bp G-rich sequence, 3’D-RS includes: 3’D-Heptamer+3’D-Spacers(23bp)+3’D-Nonamer. (3) J gene consists of 2 parts: J-RS+J-region, J-RS includes: J-Nonamer+J-Spacer(12bp)+J-Heptamer. There are 2 conservative components in J-region: [W,F]-[G,A]-X-G motif and Splicing-splice. (4) The composition of the C gene is different due to loci: TRAC and TRDC usually consist of 4 exons: Exon1+Exon2+Exon3+Exon4UTR. TRGC usually consists of 3 exons: Exon1+Exon2+Exon3, and TRGC-Exon2 usually has multiple situations: Ex2A, Ex2B, Ex2C, Ex2R, Ex2T. TRBC usually consists of 4 exons: Exon1 +Exon2+Exon3+Exon4. The fourth step is the functional identification of germline genes. The following rules are used for functional identification of germline genes: (1) 3’RSS sequence of V gene (7-23-9), J gene 5’ end RSS sequence (9-12-7), 5’ end RSS sequence (9-12-7) and 3’ end RSS sequence (7-23-9) of D gene, (2) Whether the 5’end of V gene contains leader sequence; (3) Conserved acceptor and donor splicing site; (4) Length of coding region; (5) Frameshift or stop codon of coding region. The last step is classification of group and subgroup of germline genes. For each identified and analyzed V gene, according to its existing species, a phylogenetic evolution tree is constructed to name the V gene uniformly. If no homologous V gene is found, it will be named according to its position in the loci. Each D, J and C gene was named according to its physical location in its own cluster.

### 2.2 Comparison of the TR locus of Rhinolophus ferrumequinum and other mammals

The analysis results of TRA/D, TRG and TRB loci were drawn into physical maps by Geneious Prime software according to the annotation of IMGT, and the information of four TR loci of each species in IMGT database was counted and compared. At the same time, the number of germline genes and subgroups among species were compared, and the nucleotide and amino acid composition of annotated genes were analyzed. According to the classical RSS sequence, the mutation number was counted, and the conservative analysis of 12/23 RSS sequences of V and J genes was carried out.

## 3. Results

### 3.1 TRA/TRD locus

Sequences we analyzed include: 847Kb between OR10G2 and DAD1, 174Kb between AMPH and STARD3L, and 240Kb between MOXD2 and EPHB6. Using the whole genome assembly (GCA_004115265.3) of Rhinolophus ferrumequinum recorded in the NCBI, the TRA/D locus of Rhinolophus ferrumequinum was identified (**Figure.2**): The TRA/D locus of Rhinolophus ferrumequinum was forward orientation which located on chromosome 6 (NC_046289); 5’ end borne gene-OR10G2 (2449106…2450211, GeneID: 117023609); 3’ end borne gene-DAD1 (3318653…3336546, GeneID: 117024015); The total length of the TRA locus from the first TRAV gene at the 5’end to the last TRAC gene at 3’end is about 850Kb (2456031…330546), including 81 TRAV genes (classified into 34 groups), 60 TRAJ gene (classified into 60 groups), 1 TRAC gene. TRA and the TRD locus share a part of the V gene, but the TRD locus has its own D gene, J gene, and C gene. The total length of the TRD locus from the first TRDV gene to the last reversed TRDV gene is about 660Kb (2571156…3228878), including: 18 TRDV genes (classified into 11 families), 4 TRDJ genes (classified into 4 families), 1 TRDD gene, and 1 TRDC gene. The entire organization of the TRA/D locus is: Vα(55)-Vα/δ(10)-Vδ(1)-Vα(8)-Vδ(1)-Vα(2)-Vδ(1)-Vα(4)-Vδ(2)-Dδ(1)-Jδ(4)-Cδ(1)-Vδ(1)-Jα(60)-Cα(1). The structure of TRA/D loci in bats suggests that many TRAV genes and TRDV genes are located upstream of the loci, TRD loci share a part of V genes with TRA loci, and TRD loci have a complete D-J-C cluster, including 1 TRDD gene, 4 TRDJ genes and 1 TRDC gene, and there is a TRDV gene with opposite transcription direction downstream of TRDC gene, and finally a TRA locus J-C cluster, including 60 TRAJ genes and 1 TRAC gene. **Supplementary Table 1** provides detailed information for each annotated TRA/D locus germline genes in Chromosome6.

**Figure.2.**
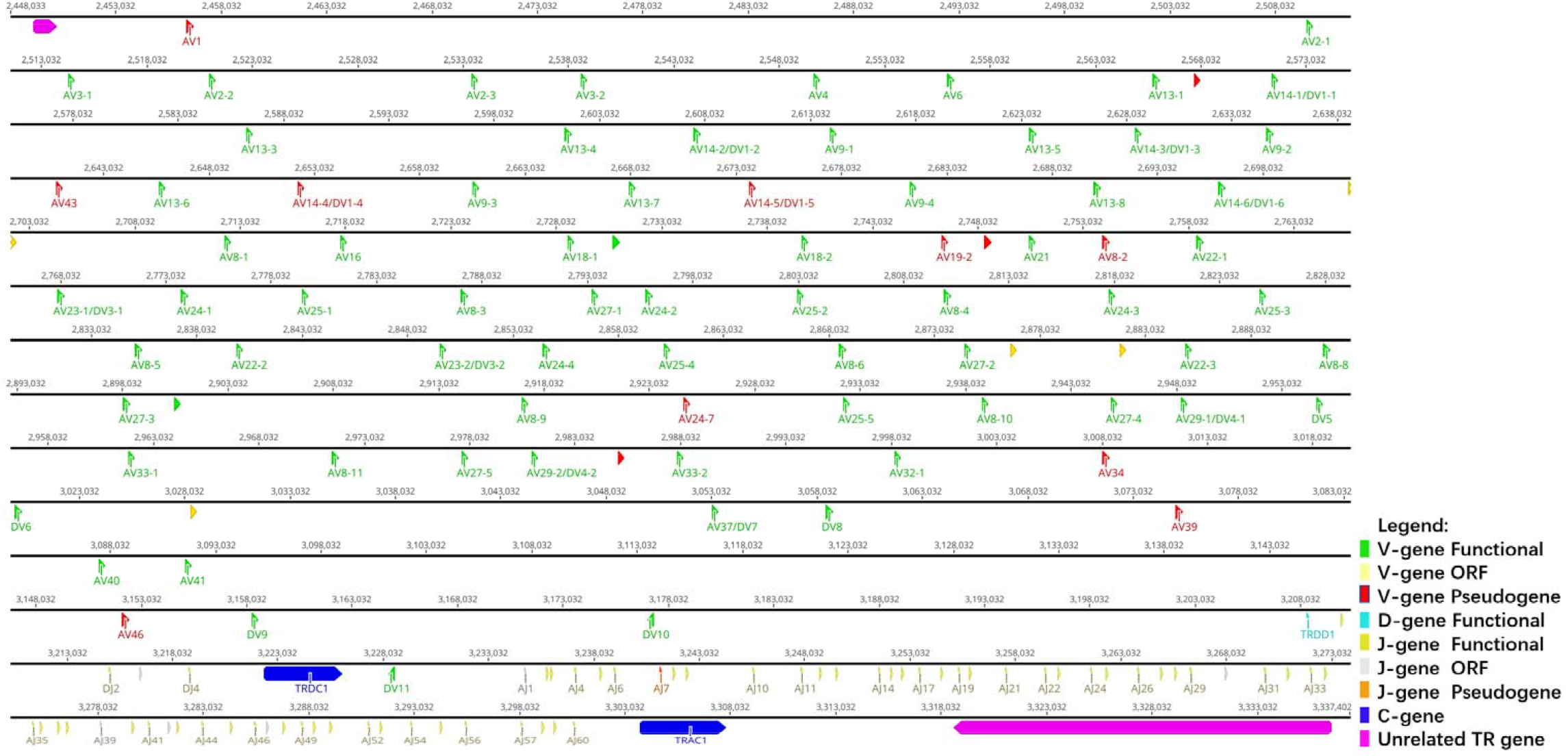
The schematic representation of the genomic organization of the bat TRA/D locus. The diagram shows the position of all the related and unrelated TRA/D genes according to nomenclature. The L-part and RSSs are not shown. The arrows indicate the transcriptional orientation of the genes. The number represents the specific location of the genes in the chromosome 6 of Rhinolophus ferrumequinum (GCA_004115265.3)

### 3.2 TRG locus

The TRG locus (**Figure.3**) of the (Figure.3) is located on chromosome 20 (NC_046303) in reverse orientation: 5’ end borne gene-AMPH (52427198…52205147, GeneID: 117012456); 3’ end borne gene-STARD3NL (52030211…51976009, GeneID: 117012446); The whole length of TRG locus from the first TRGV gene at 5’ end to the last TRGC gene at 3’ end is about 150Kb(52427198…51976009), including 14 TRGV genes (classified into 7 families), 6 TRGJ genes (classified into 3 families) and 2 TRGC (classified into 2 families) genes. The organization of TRG locus is: Vγ(7)-Jγ(4)-Cγ(1)-Vγ(7)-Jγ(4)-Cγ(1). It suggests that the TRG locus of Rhinolophus ferrumequinum is composed of two V-J-C gene clusters, and each gene cluster is composed of the same number of genes, including 7 V genes, 4 J genes, and 2 C gene. **Supplementary Table 2** provides detailed information for each annotated TRG locus germline genes in Chromosome20.

**Figure.3.**
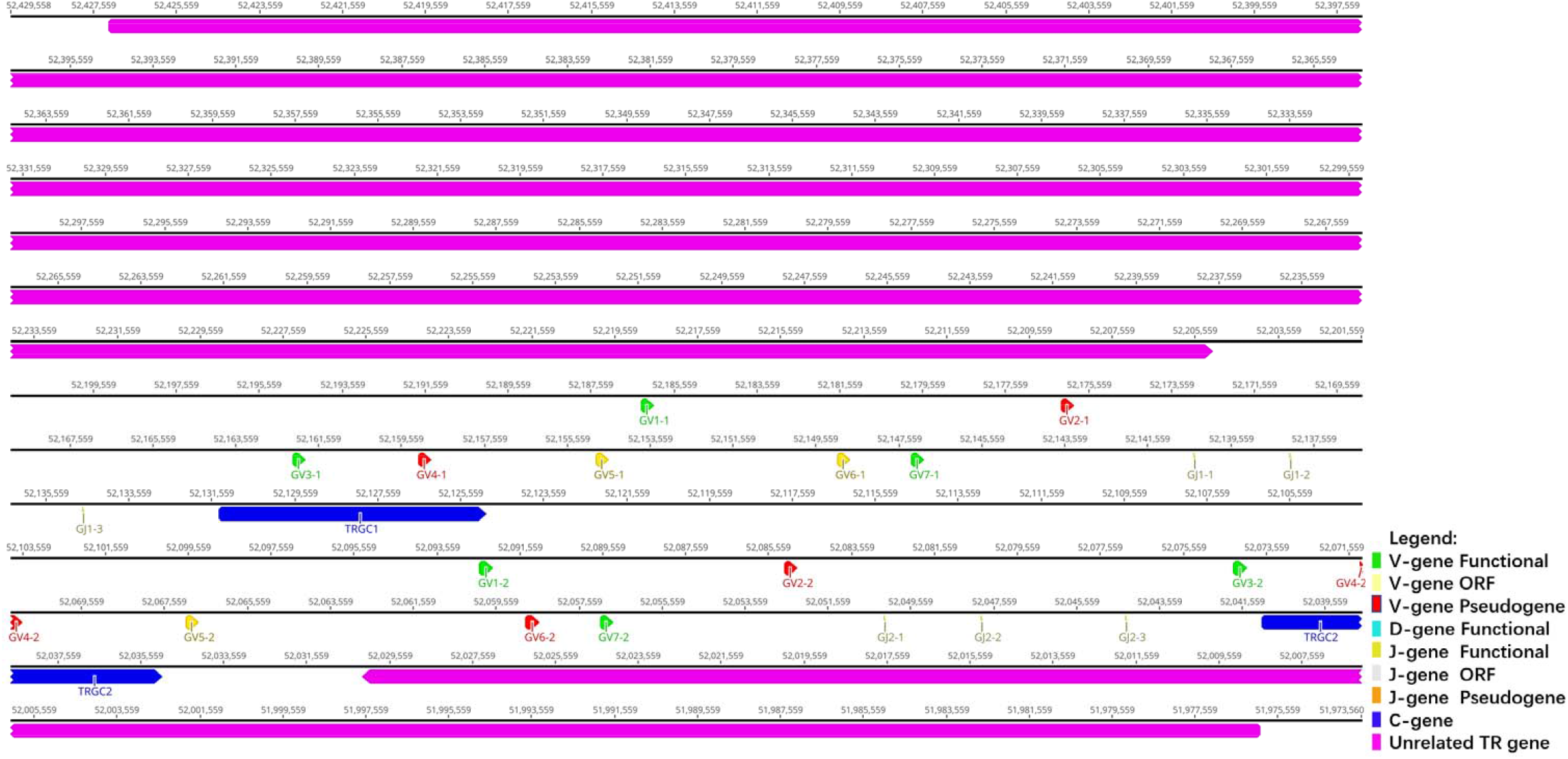
The schematic representation of the genomic organization of the bat TRG locus. The diagram shows the position of all the related and unrelated TRG genes according to nomenclature. The L-part and RSSs are not shown. The arrows indicate the transcriptional orientation of the genes. The number represents the specific location of the genes in the chromosome 20 of Rhinolophus ferrumequinum (GCA_004115265.3)

### 3.3 TRB locus

The TRB locus (**Figure.4**) is located on chromosome 26 (NC_046309) in reverse orientation: the 5’ end borne gene-MOXD2 (8164121… 8171343, Gene ID: 117018015); The 3’ end borne gene-EPHB6 (7911840… 7926436, Gene ID: 117018107); The total length of TRB locus from the first TRBV gene at 5’ end to the last reverse TRBV gene at 3’ end is about 203Kb(8159483…7911840), including 29 TRBV genes (classified into 25 families), 2 TRBD genes (classified into 2 families), 15 TRBJ genes (classified into 2 families) and 2 TRBC genes. The entire organization of TRB locus is: Vβ(28)-Dβ(1)-Jβ(6)-Cβ(1)-Dβ(1)-Jβ(9)-Cβ(1)-Vβ(1).

**Figure.4.**
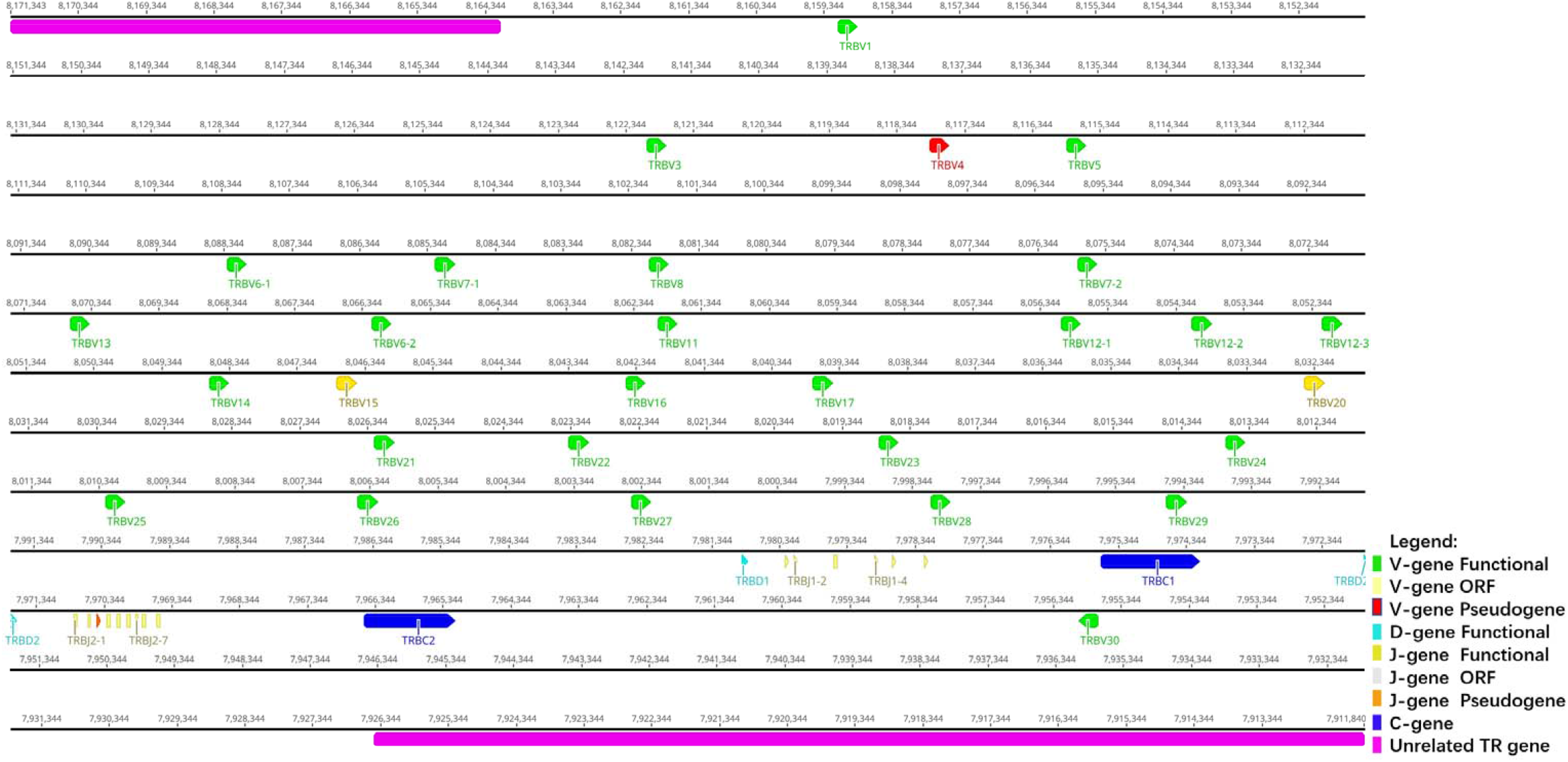
The schematic representation of the genomic organization of the bat TRB locus. The diagram shows the position of all the related and unrelated TRB genes according to nomenclature. The L-part and RSSs are not shown. The arrows indicate the transcriptional orientation of the genes. The number represents the specific location of the genes in the chromosome 26 of Rhinolophus ferrumequinum (GCA_004115265.3)

The structure of the TRB locus of Rhinolophus ferrumequinum suggests that the upstream is a V cluster composed of 28 TRBV genes, and the downstream is two complete D-J-C gene clusters. The first cluster contains: 1 D gene, 6 J genes and 1 C gene, the second cluster contains: 1 D gene, 9 J genes and 1 C gene. There is a TRBV gene with the opposite transcription direction downstream of the second TRBC gene. **Supplementary Table 3** provides detailed information for each annotated TRB locus germline genes in Chromosome26.

### 3.4 Classification of germline gene in 4 TR locus of Rhinolophus ferrumequinum

#### 3.4.1 Classification and phylogenetic analysis of the V genes

(1) A total of 81 TRAV genes were found in TRA loci, and the nucleotide similarity of 81 TRAV genes ranged from 25.1% to 98.9%. According to the standard that the nucleotide homology is more than 75% and belongs to the same group, it can be divided into 31 groups.

In order to analysis the classification and affiliation between annotated V genes of bat with existing species, we selected the V gene sequences of primate representative species, human, carnivorous representative species, dog and artiodactyl representative species, cattle, from the IMGT database to construct phylogenetic trees. There are two criteria for selecting genes: one is to select only potential functional genes and in-frame pseudogenes, and the other is to select only one gene for each subgroup.

After all the selected V genes were compared by ClustalW method, MEGA7 was used for N-J method of rootless tree construction. The phylogenetic tree constructed by 31 bat TRAV genes shown in **Figure5-A**. The results reflect the homology between the annotated bat TRAV genes and existing species. We named the bat TRAV genes according to the comparison results.

**Figure.5.**
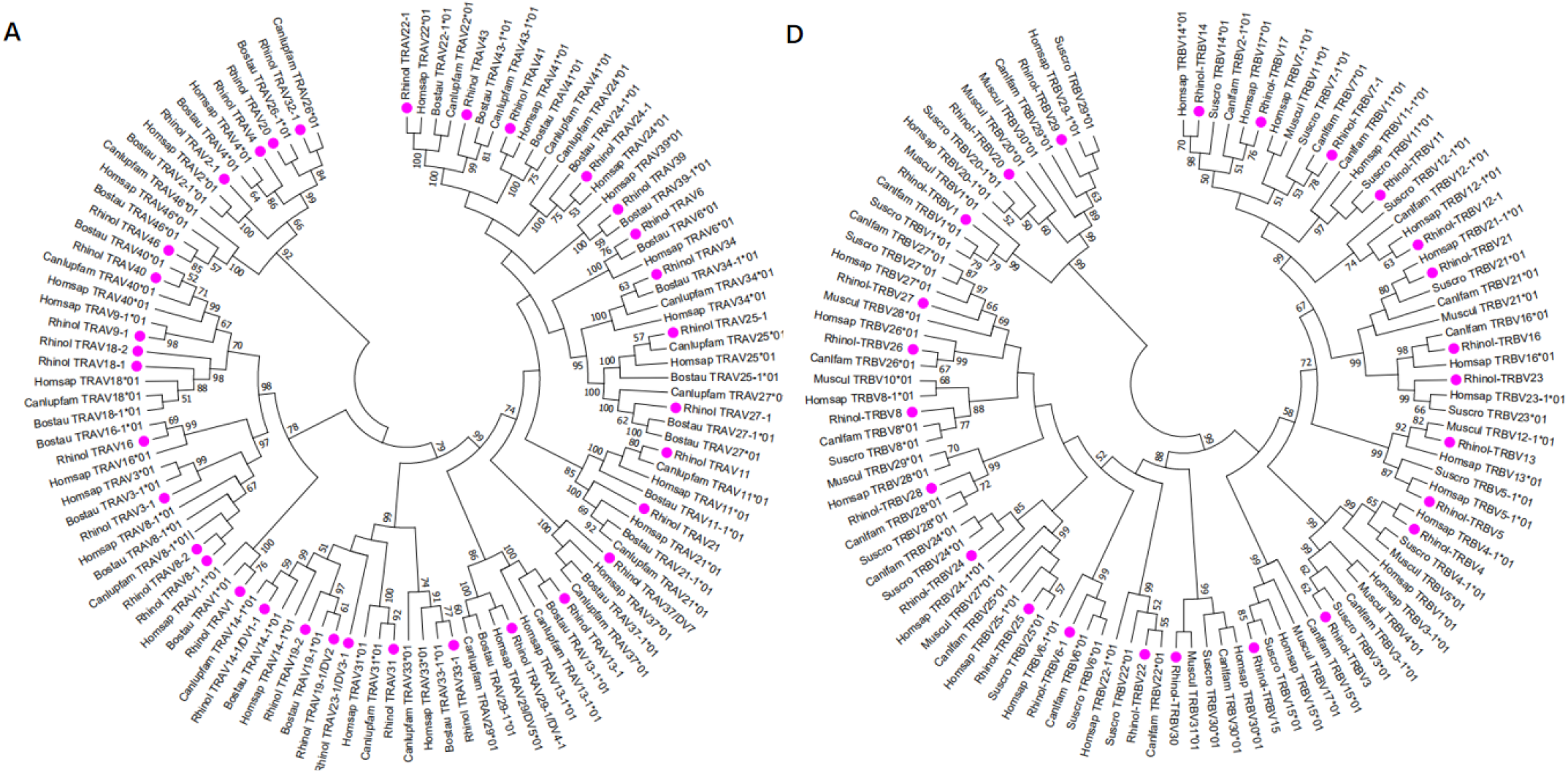
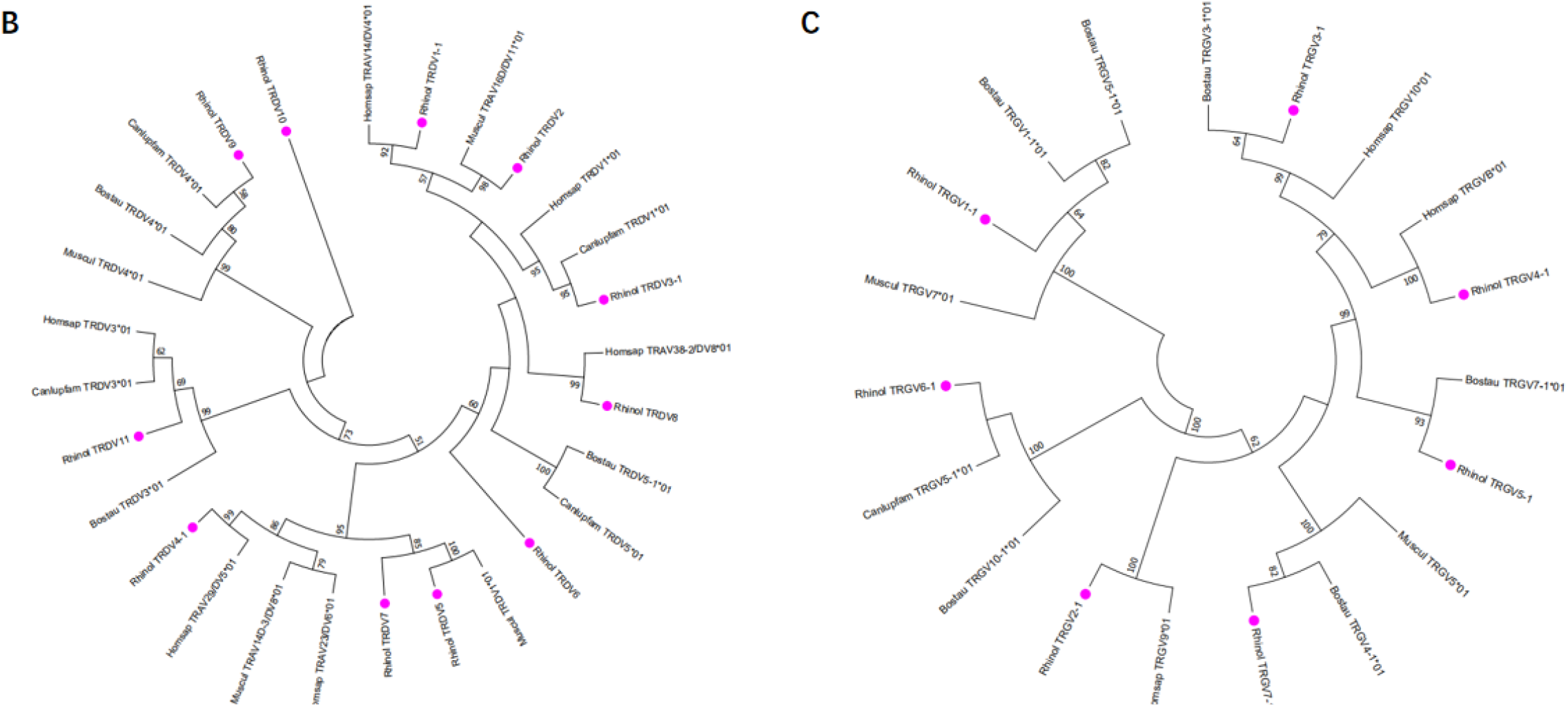
Phylogenetic analysis of TRAV(A)、 TRDV(B)、 TRGV(C)、 TRBV(D)genes. Unrooted trees were constructed using the Neighbor-joining method based on V-REGION nucleotides sequences of Bat、 Dog、 Human、 Mouse、 Cow. The percentage of the nodes in 1000 bootstrap replicates are show on the branches. Bat genes are labelled in pink. The IMGT standardized abbreviation for taxon is used: six letters for species (Rhinol, Homsap, Bostau, Muscul) and nine letters for subspecies (Canlupfam)

It should be noted that there are 3 groups, TRAV8, TRAV18 and TRAV19, of which the TRAV8-2 gene and the rest of the TRAV8 members are only about 65% identity, but in the evolutionary branch, we found that TRAV8-2 and TRAV8-1 clustered to the same branch, since TRAV8-2 is a pseudogene, we incorporated it into TRAV8 group. The same situation occurs when the identity between TRAV18-1 and TRAV18-2 is 73.8%, and TRAV19-1 and TRAV19-2 is 69.6 %. Of the 31 families, 27 families can form branches with human and cattle genes at the same time. Genes that failed to find homology are named according to their positions in the locus. There are 16 multigene groups in TRAV, and 6 groups have been significantly amplified, including TRAV8, TRAV13, TRAV14, TRAV24, TRAV25, and TRAV27 (each family has 5 members), and 10 groups are composed of 2-4 members. Including TRAV2, TRAV3, TRAV9, TRAV18, TRAV19, TRAV22, TRAV23, TRAV29, TRAV32, TRAV33, and the remaining 20 are single gene group.

(2) There are 18 TRDV genes in the TRD locus, and the nucleotide identity is between 27%-98.6%. The 18 TRDV genes are divided into 11 families, TRDV5, TRDV6, TRDV7, TRDV8, TRDV9, TRDV10, TRDV11 are single gene groups. The phylogenetic tree constructed by the 11 TRDV genes of bat with human, dog, and cattle is shown in **Figure 5-B**. Except for the TRDV10 family, all groups of the bat can find homology in the 3 species. Of the 18 TRDV genes, only 2 shared with TRA loci were classified as pseudogenes, and the other 16 were functional genes.

(3) The TRG locus contains 14 TRGV genes with nucleotide similarity ranging from 33% to 99%. Because the TRG locus of bat is composed of two identical V-J-C gene clusters, the 14 TRGV genes are divided into 7 groups with nucleotide identity ranging from 94.6% to 99.0%. A phylogenetic tree was constructed with the annotated TRDV genes of bat and the TRDV genes of human, dog, cattle, and mice (**Figure5-C**). All TRGV genes of bat can be found homologous in 4 species. The 14 TRGV genes have a high proportion of pseudogenes, 5 of 14 are classified as pseudogenes, 3 ORFs and 6 functional genes. Because the recorded data of TRDV and TRGV genes between species are limited and the uniformity is not high, they are named according to position in the locus.

(4) The TRB locus contains 29 TRBV genes with nucleotide identity ranging from 33.7% to 94.2%, and the 29 TRBV genes are divided into 25 groups. The phylogenetic trees constructed by human, mouse, dog, and pig showed that all the annotated TRBV genes of the bat could be found to be homologous in 4 species. It is worth noting that human TRBV1 could not cluster with the TRBV1 group of cattle, mouse, pig and bat, but was on the branch of TRBV4. There are 3 multigene groups, TRBV5 and TRBV6 have 2 gene members, and TRBV12 has 3 gene members. The 29 TRBV genes have 1 pseudogene and 2 ORFs, and the remaining 26 are classified as functional genes. The descriptions of pseudogenes and ORFs of all germline genes of TR locus of bat are shown in **Supplementary Table 4**.

The amino acid sequences of some TRAV genes (81 TRAV genes in **Supplementary Figure 1**) and all TRDV, TRGV and TRBV genes annotated of bat were aligned according to the unique number of IMGT V-region (**Figure 6**), so as to ensure the homology to the maximum extent.

**Figure.6.**
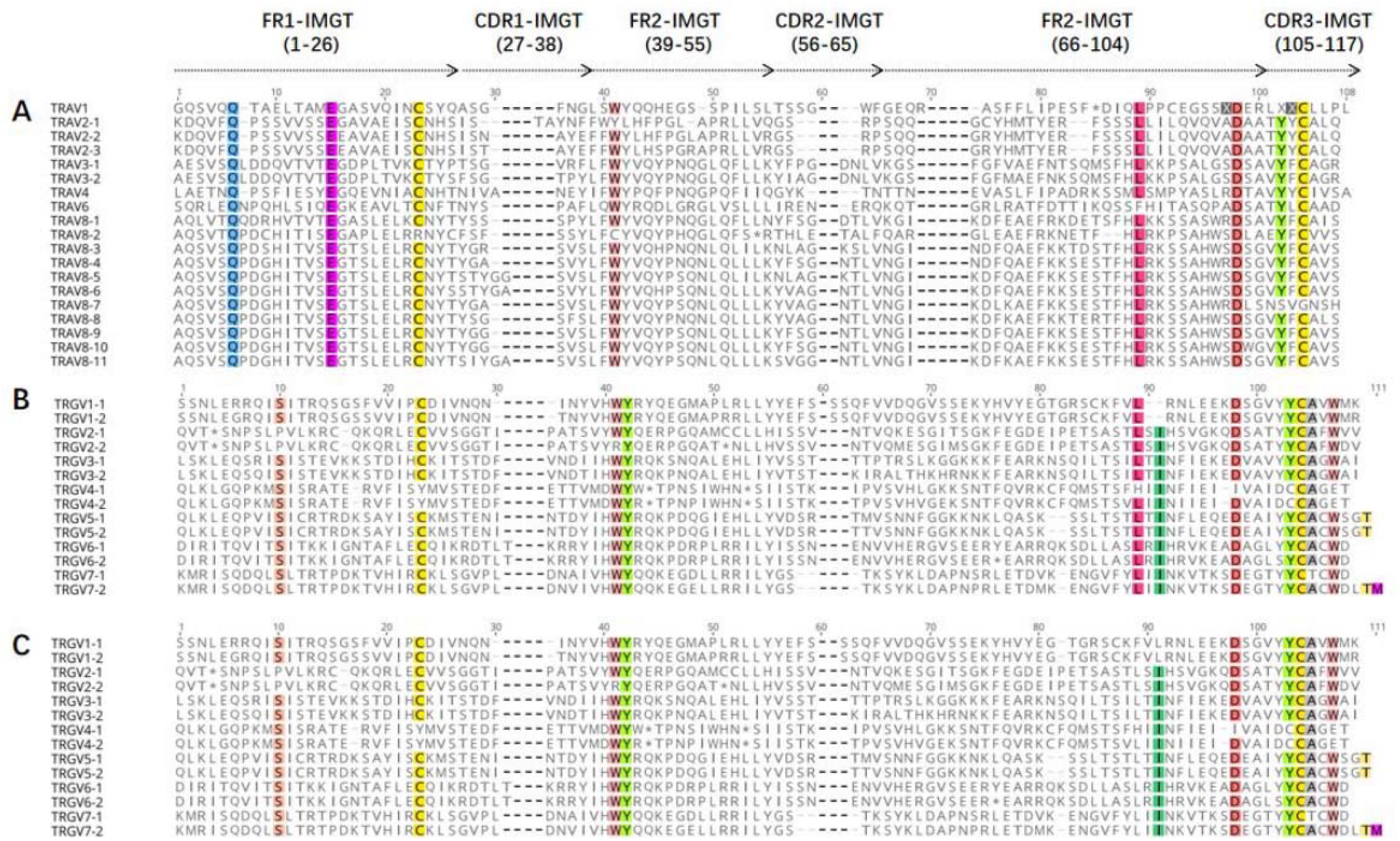
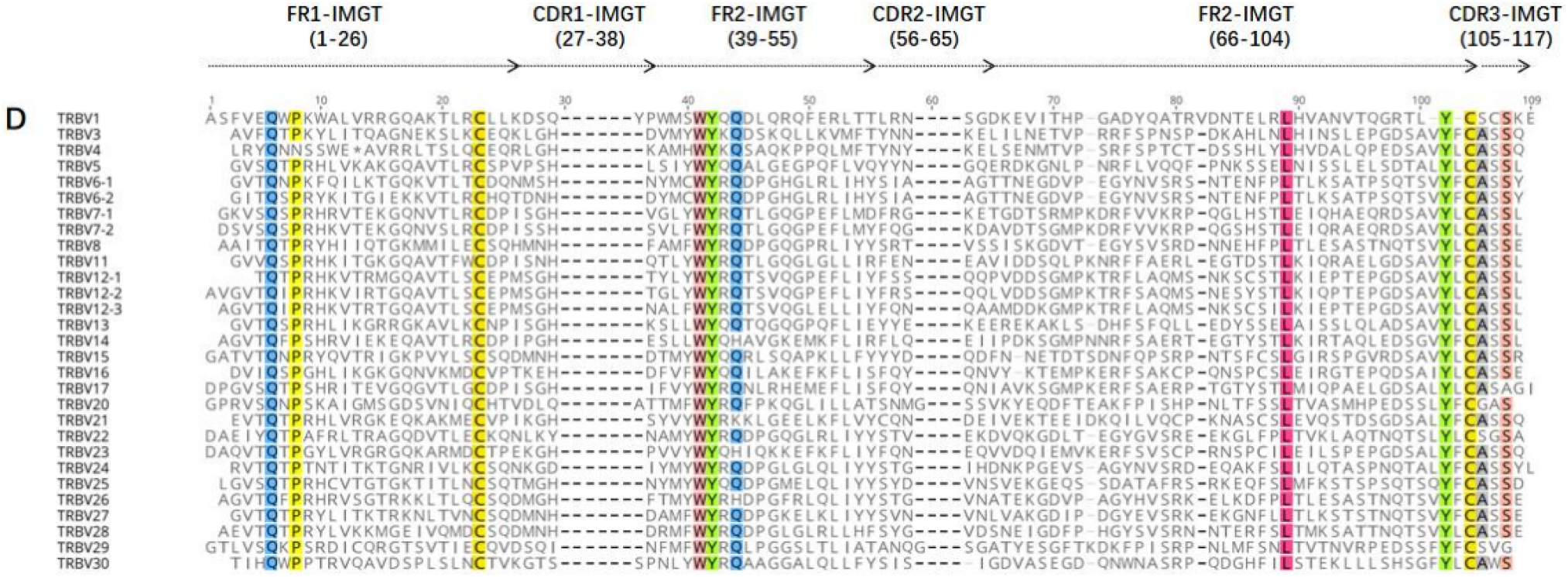
The IMGT Protein display of the bat TRAV(A), TRDV(B), TRGV(C), TRBV(D) genes. TRAV genes are only shown TRAV1 to TRAV10. The description of the FR-IMGT and CDR-IMGT is according to IMGT unique numbering for V-region. The four conserved amino acid of the V-domain (1st-CYS 23, CONSERVED-TRP 41, hydrophobic AA 89,2nd-CYS 104) are indicated in color

#### 3.4.2 Analysis of D gene and J gene

According to the naming rules stipulated by IMGT, J genes are classified by C genes and numbered according to the positions in their respective (D)-J-C gene clusters. The length of J gene is 51-66bp, and the typical [W, F]-[G, A]-X-G motif is retained, which determines the function of J gene. There are 12RSS sequences at the 5’ end and donor splicing sites at the 3’ end of each J gene.

We have annotated a total of 60 TRAJ genes with nucleotide similarity between 22.2% and 71.9%. According to nucleotide homology, they can be divided into 60 families. The nucleotide and amino acid deduced sequences of TRAJ are shown in **Fig7-A** (Supplementary Figure 2 for 60 TRAJs). Only TRAJ1 and TRAJ7 do not contain the conservative [W, F]-[G, A]-XG motif. Among the 60 TRAJ genes, TRAJ1, TRAJ30, TRAJ39, TRAJ42, and TRAJ47 are classified as ORF, TRAJ7 is classified as pseudogene, and the remaining 54 TRAJ are functional genes; A total of 4 TRDJ genes are annotated with nucleotide similarity between 48.9%-66.7%, which can be divided into 4 families. The nucleotide and amino acid deduced sequences of TRDJ are shown in **Figure 7-B**. 3 TRDJ genes contain conserved FGXG motif is therefore classified as a functional gene, only TRDJ3 is classified as an ORF; a total of 6 TRGJ genes are annotated with nucleotide similarity between 53.1% and 100%, which can be divided into 2 families. The acid and amino acid deduced sequence is shown in **Figure 7-C**. The similarity between TRGJ1-1 and TRGJ2-1 is 100%, TRGJ1-2 and TRGJ2-2 is 98.3%, TRGJ1-3 and TRGJ2-3 is 98%, the 6 TRGJ genes all contain conserved F- [G, A]-XG motif and RSS sequences, so they are classified as functional genes. A total of 15 TRBJ genes were annotated. Since the TRB locus contains 2 D-J-C clusters, the 15 TRBJ genes are divided into two families, and their nucleotide similarity ranges from 33.3% to 91.1%. the similarity of TRBJ-4 with TRBJ-5 and TRBJ-6 reaches 91.1% and 84.4%, respectively. The nucleotide and amino acid deduced sequences are shown in **Figure7-D**. Only 1 D gene was found in the TRD locus. The nucleotide and deduced AA sequence of TRDD1 gene are shown in **Figure7-E**, which consists of a 14bp G-rich fragment, and the RSS sequences of upstream and downstream are very conservative. Two D genes were found in TRBV locus. the nucleotide and deduced AA sequences of TRBD1 and TRBD2 genes are shown in **Figure7-E**. The two TRBD genes are composed of 12bp and 15bp G-rich fragments respectively.

**Figure.7.**
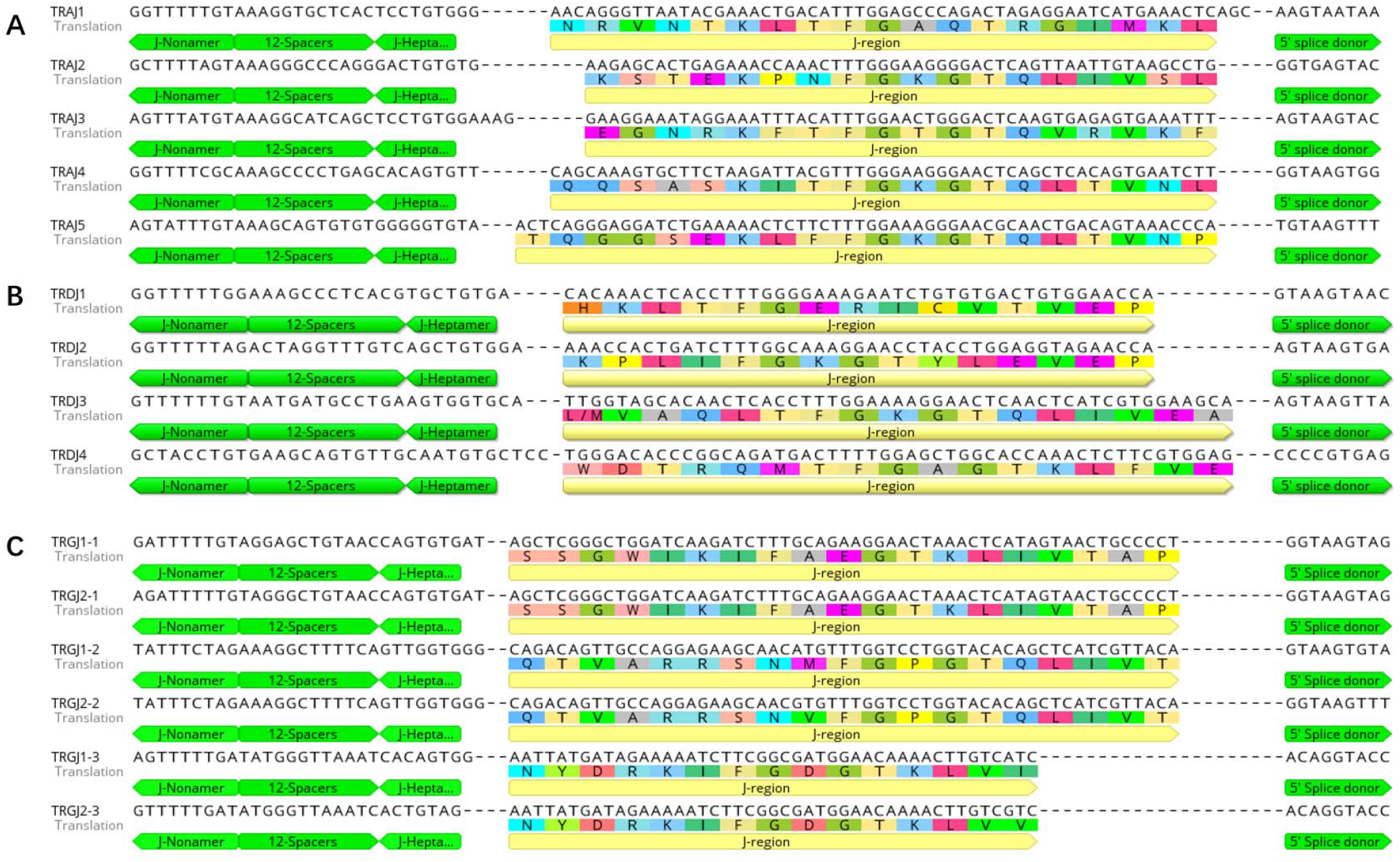
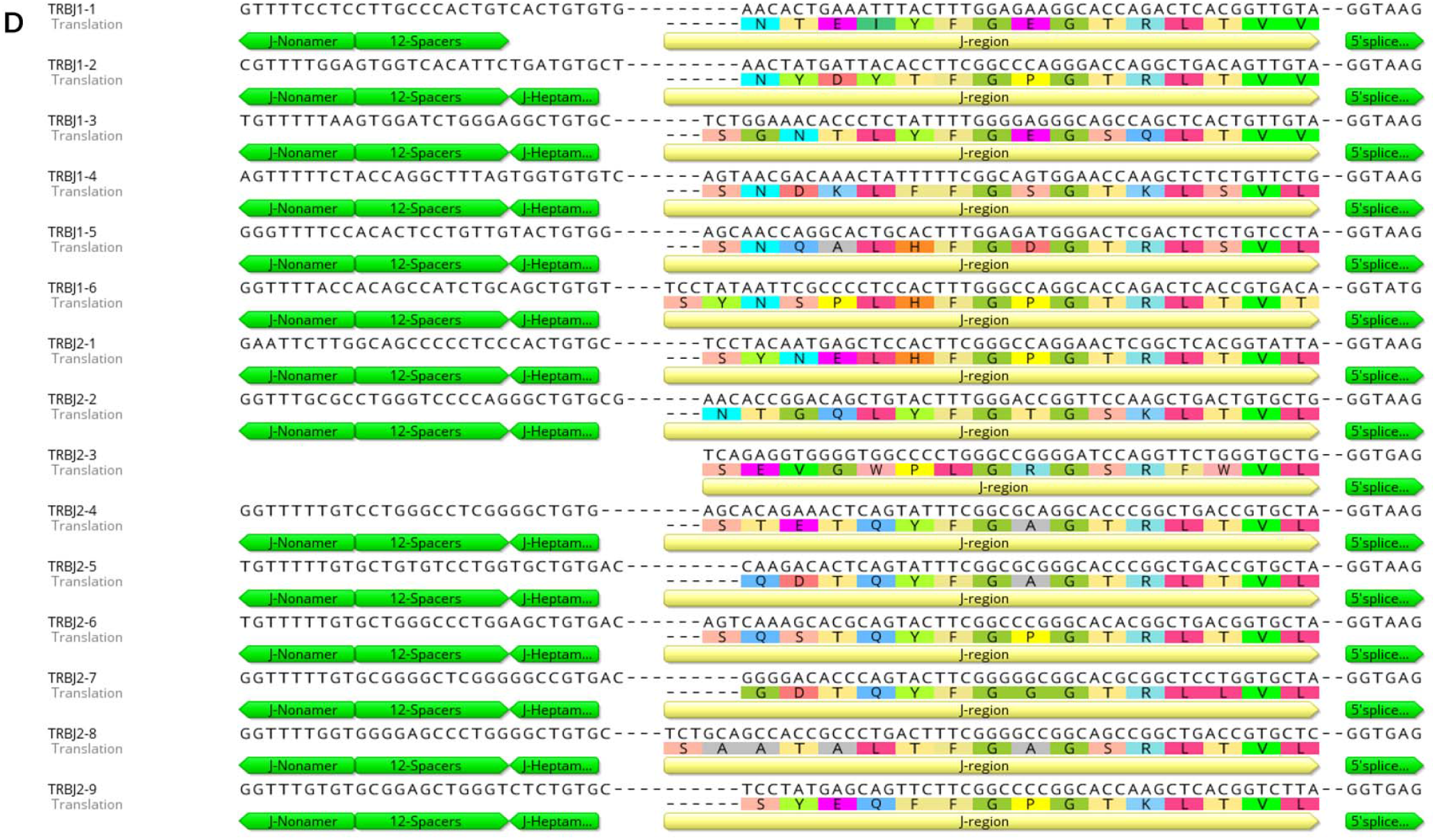
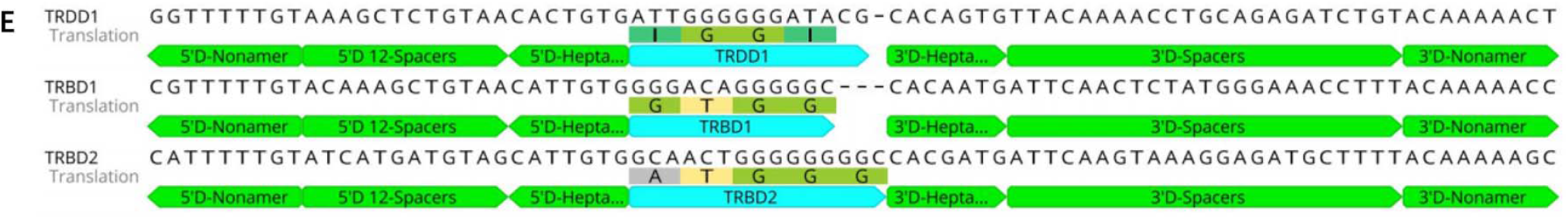
Nucleotide and deduced amino acid sequences of the bat TRAJ(A)、 TRDJ(B)、 TRGJ(C)、 TRBJ(D)、 TRBD and TRBD(E) genes. The heptamer、 nonamer and splice donor is labeled. The numbering adopted for the gene classification is reported on the left of each gene

#### 3.4.3 Analysis of C gene

At the genome level, each C gene is composed of multiple exons, EX1 encodes a constant region, part of the 5’ ends of EX2 and ex3 encodes a connecting region, part of the 3’ end of ex3 and the first codon of EX4 encodes a transmembrane region, and the rest of EX4 encodes a cytoplasmic region. The fourth exon of TRAC and TRDC is usually untranslated Exon4UTR, while TRGC does not have the fourth exon, only TRBC usually contains a complete amino acid sequence of the four exons. Exon1, Exon2 and Exon3 have the same size, but different species have different introns. We analyzed the amino acid sequence and intron/exon structure of the annotated C gene of bat with each species (**Figure 8 and Supplementary Figure 3**). We found 1 TRAC gene and 1 TRDC gene in the TRA/D locus, respectively. The TRAC1 gene of bats encodes 136 amino acids. The amino acid similarity of the TRAC gene between species is between 47.1% and 81.4%. The highest similarity between TRAC gene of bat and rabbit is 54.5%. Exon1 of TRAC varies from species to species, while Exon2 and Exon3 are highly conserved. The TRDC1 gene of bat encodes 142 amino acids, and the amino acid sequence similarity of the TRDC gene of each species ranges from 64.1% to 86.35. The similarity between the TRDC gene of bat and rhesus monkeys reaches 72.6%, except for Exon4UTR, the remaining three exons are highly conserved. There are two TRGC genes in the TRG locus of the bat, both of which are encoded by 163 amino acids. Because TRGC has multiple Exon2 structures, the intron length and structure of TRGC genes in different species are varied. However, we only matched one Exon2 among the two TRGC genes of the bat. There are two TRBC genes in the TRB locus of the bat, both of which are encoded by 175 amino acids and consist of 4 complete exons. The similarity of amino acid sequences among TRBC1 species is between 72.5% and 92.6%, and the similarity of TRBC2 is between 71.3% and 93.8%.

**Figure.8.**
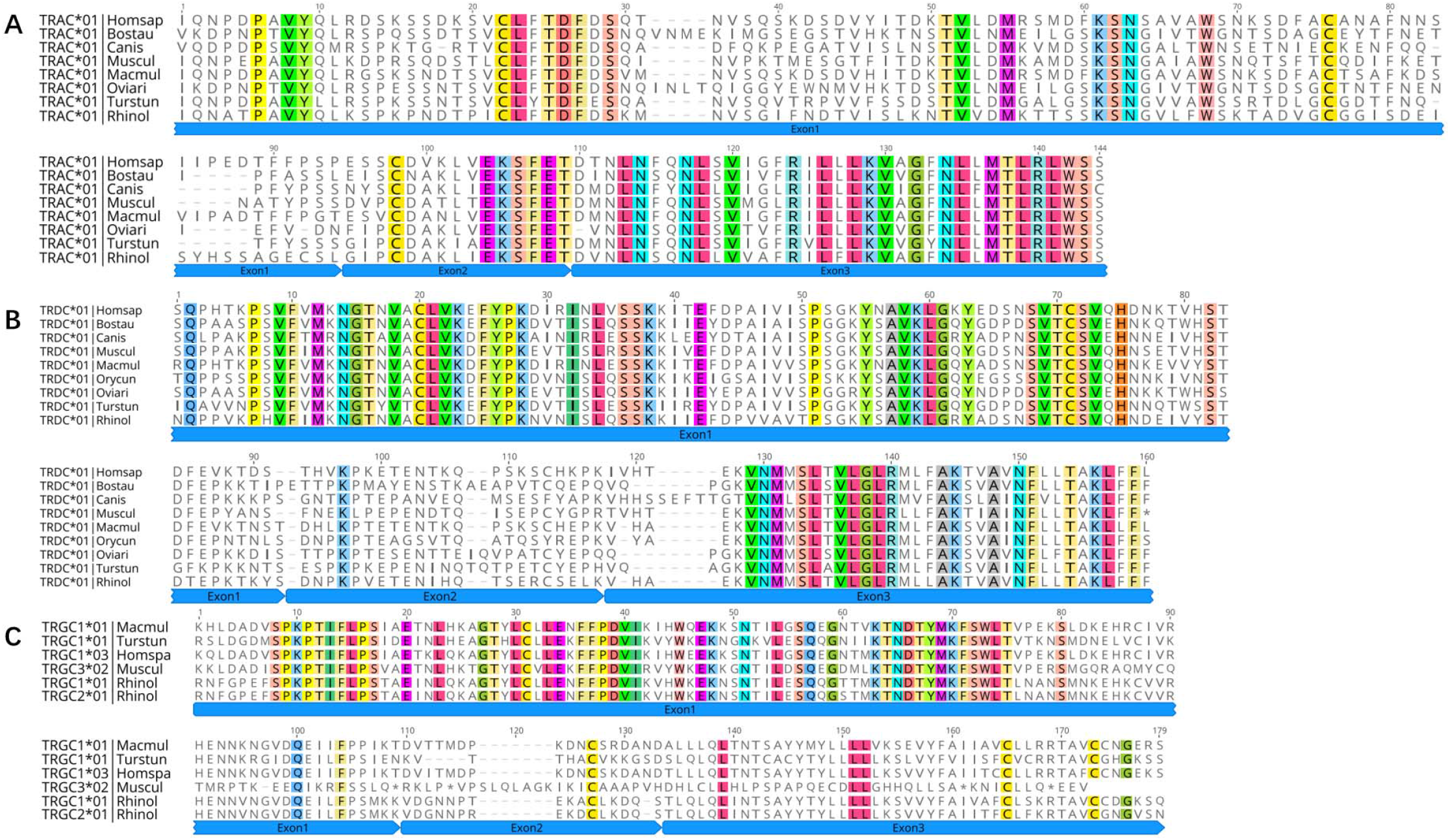
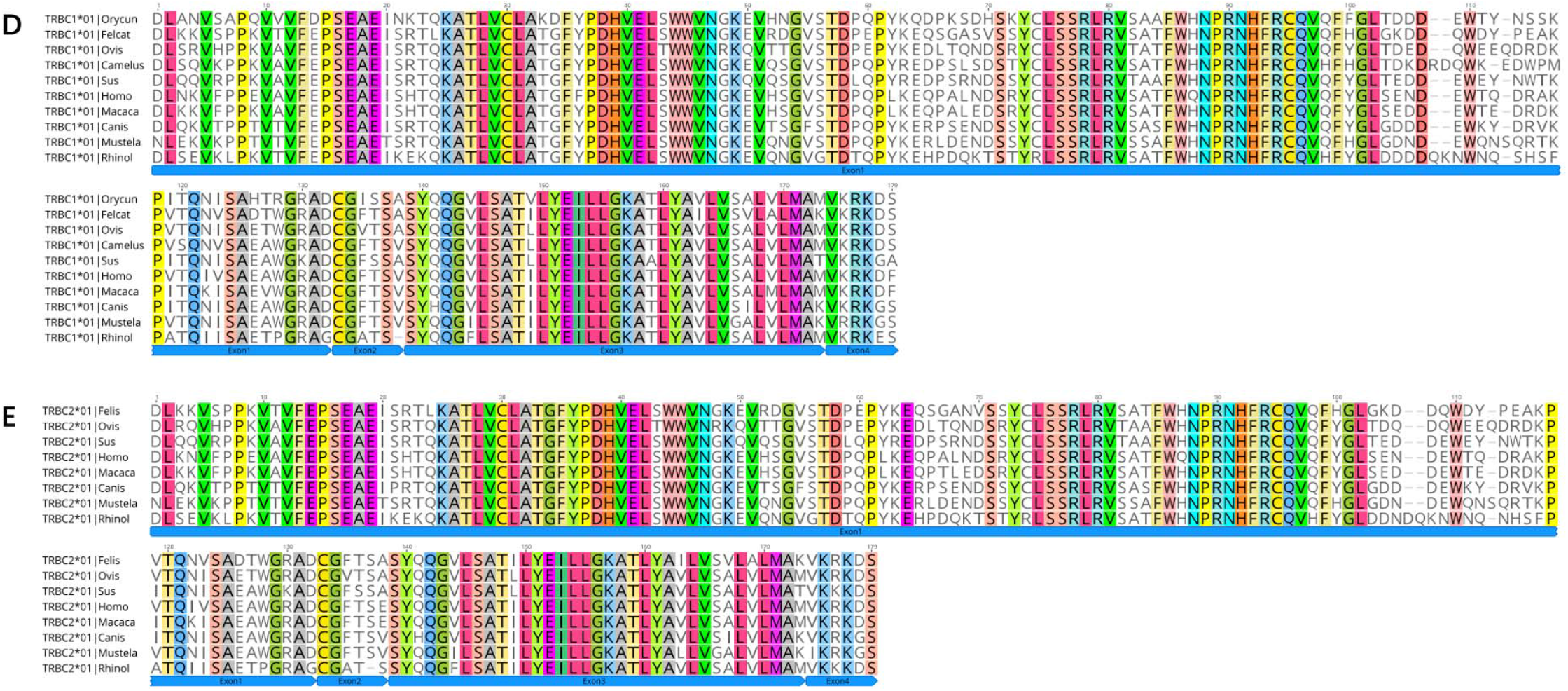
Deduced amino acid sequences of the bat TRAC(A)、 TRDC(B)、 TRGC(C)、 TRBC1(D) and TRBC2 genes. Felcat: cat, Canlupfan: dog, Bostau: cow, Oviari: sheep, Turstun: dolphin, Orycun: rabbit, Macmul: rhesus monkey, Homspa: human, Muscul: mouse, Rhinol: greater horseshoe bat

### 3.5 Germline gene recombination signal sequences analysis

According to the IMGT rule, we statistically analyzed the RSS sequences judged as germline genes (**Figure 9**). We used the classic 7-mer-CACAGTG (heptamer) and 9-mer-ACAAAAACC (nonamer) as motifs, The number of mutations in the RSS sequence was analyzed: In the TRA/D locus, 7 of the 81 TRAV genes did not have the 23RSS sequence, and the number of mutations in the 74 RSS sequences was between 0-9; The 3’ ends of 18 TRDV genes all have 23RSS sequences, and the number of mutations in 18 RSS sequences is between 0 and 9, the number of mutations in 15 sequences is less than 6, the number of mutations in 2 sequences is 8, and the number of mutations in one sequence is 9. Of the 60 TRAJ genes, 5 TRAJ have no 12RSS sequence at the 3’ end, and the number of mutations in 55 12RSS sequences is between 0 and 7, among which the number of mutations in 54 sequences is less than 6 and the number of mutations in 1 sequence is 7. In TRG locus, 2 out of 14 TRGV genes do not have 23RSS sequence, and the number of mutations in 23RSS sequence of 12 TRGV genes is between 0 and 9, of which 11 have less than 7 mutations, and the number of mutations in one sequence is 9. The 6 TRGJ genes all have 12RSS sequences at the 3’ end, and the number of mutations in the 12RSS sequences of the 6 TRGJ genes ranges from 0 to 9, among which the number of mutations in 3 genes is less than 5 and the number of mutations in 3 genes is 9. In TRB loci, 29 TRBV genes all have 23RSS sequences, and the number of mutations in 23RSS sequences of TRBV genes is between 1 and 6, of which only 1 mutation number is 6, and a total of 20 mutations are concentrated in 3-4 mutations; Only 1 of the 15 TRBJ genes does not have 12RSS sequence, and the number of mutations in 12RSS of 14 genes is between 2 and 6, mainly between 3 and 5. It can be seen that the RSS sequence of TRB locus is more conservative than other loci.

**Figure.9.**
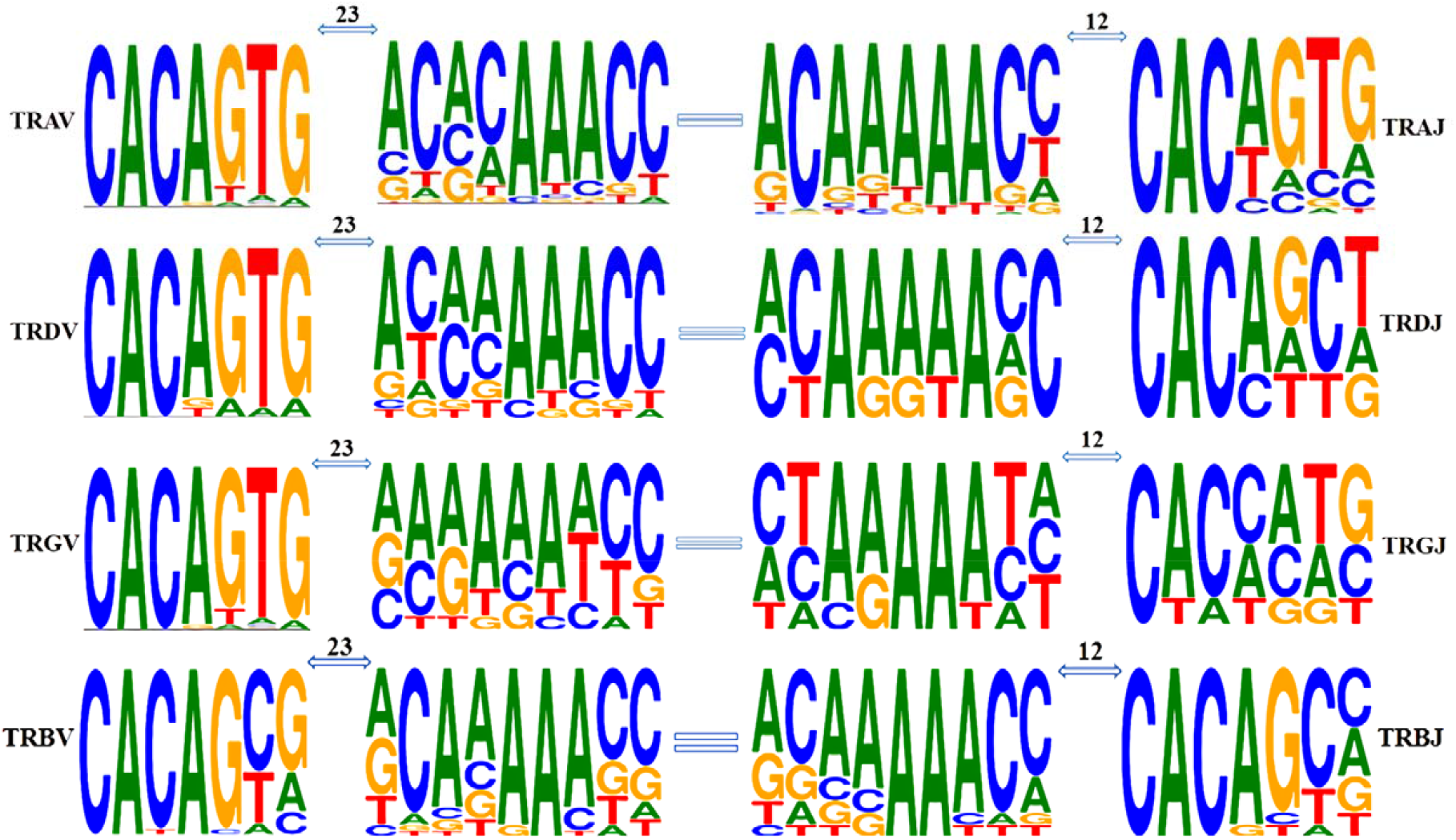
Position weight matrixes of recombination signal sequences of T cell receptor V and J genes. The height of symbols indicates the relative frequency of each nucleotide at that position

All the V genes heptamer and the poly-A tract of V gene nonamer are highly conserved. The nonamer of TRAV and TRDV has a high degree of diversity in the first 4 nucleotides, while the whole nonamer of TRGV has high diversity. The diversity of RSS sequence is more obvious in J gene, except that the poly-A tract of J gene nonamer is conservative. The first 2 nucleotides and the last two of nonamer of J gene are diversified, which is more obvious in heptamer of J gene, except the bases (cac) of the first 3 nucleotides, the last 4 nucleotides of J gene heptamer are highly diversified.

## 4. Discussion

The function of vertebrate T/B cells depends on TCR/BCR, and the TR locus analysis of species is of great significance for us to understand its immune response. The information of 4 TR loci of many species has been included in the IMGT database, and the TR loci of 12 species, including human, macaque, mouse, pig, cow, sheep, dog, cat, dolphin, ferret, rabbit and zebra fish, have been completely annotated and described(http://www.imgt.org/IMGTrepertoire/LocusGenes/#C). Previous studies always annotated and analyzed one or two loci of a single species. However, for the first time, we annotated the 4 loci TRA/TRD, TRG and TRB of Rhinolophus ferrumequinum directly through the whole genome through the distribution characteristics, sequence characteristics and conservative RSS of germline genes. This paper can provide ideas and reference methods for new species with unresolved TR/IG loci. In addition, we believe that high-quality TR locus annotation is the key to the subsequent analysis of the TCR/BCR repertoire of Rhinolophus ferrumequinum.

Gene replication is the basis of the evolution of antigen-specific receptors, which leads to the formation of independent IG/TR sites [26, 27]. Since mammalian TRD loci are embedded in TRA loci, the length of this locus spans the longest among the TR loci, even so, the structure has not changed much [28], and the amplification events in the genome are part of the reasons for the differences. In the TRA/D locus, the most obvious replication events occurred in Artiodactyl cattle (3330Kb) and sheep (2880Kb). In terms of the structural characteristics of the loci, the TRA/D locus of Rhinolophus ferrumequinum also maintained a conservative structure of mammals, only primate macaques (806Kb), carnivorous cats (830Kb) and dogs (760Kb) are shorter than the TRA/D locus of Rhinolophus ferrumequinum (850Kb).

In contrast, although the coverage of TRG loci is the shortest, the situation among species may be more complicated. Many species have detected the existence of 2 TRG loci. For example, there are two loci, TRG1 and TRG2, in Atlantic salmon, although the expression of TRG2 cannot be detected [29]. In addition, the study of sheep found that there were at least two TRG loci on chromosome 4, and the expression analysis showed that both of them had functions [30, 31], and the second TRG locus was found in cattle, buffalo and goats by the same method [32, 33]. On the other hand, the TRG locus structure of Rhinolophus ferrumequinum is relatively simple, which consists of two V-J-C gene clusters, and the composition and similarity among the gene clusters are almost identical. The dog’s TRG locus has undergone repeated replication, resulting in eight V-J-C gene clusters [34]. Mammalian TRB locus is probably the most conservative, with multiple V genes in the upstream and multiple D-J-C clusters in the downstream, and replication events occur among D-J-C gene clusters. In the early analysis of chicken expression level, only one D-J-C cluster was found [35]. Primate humans, macaques, rodent mice, carnivorous cats, dogs, and Rhinolophus ferrumequinum all contain two D-J-C clusters. Artiodactyla is the representative of three D-J-C gene clusters, including pigs, sheep. We found that the TRBJ gene of the second D-J-C cluster and the third cluster of pigs and sheep has very high homology, so the replication event is likely to occur in the second gene cluster, and there are four D-J-C clusters downstream of the TRB locus of short-tailed opossum [36]. In addition, the TRB loci of primate humans (620Kb) and rhesus monkeys (736Kb) have the largest span, and the number of embryonic genes is only slightly less than that of sheep (506Kb). It is worth noting that a TRDV gene and a TRBV gene with opposite transcription directions are arranged in the downstream of TRDC and the downstream of TRBC in mammals, and these two reverse transcribed V genes are expressed in both human and mouse. The 81 TRAV genes of the Rhinolophus ferrumequinum can be classified into 31 subgroups, among which 14 subgroups are multigene families, and obvious amplification occurs in TRAV8(11 members), TRAV13(8 members), TRAV14(6 members) and TRAV24(7 members). Although 54 human TRAVs can be classified into 44 subgroups, there are only 7 multigene groups, and only TRAV8 has more than 3 members; The most significant is the 183 TRAV of cattle, which are classified into 42 subgroups, of which 34 are multigene groups, and TRAV22, TRAV23, TRAV25 and TRAV26 all have obvious amplification [40]. In the TRDV of 18 Rhinolophus ferrumequinum, except TRDV1(6 members), TRDV3(2 members) and TRDV4(2 members), all the other families are single gene families, and the amplification of TRA/D locus is particularly remarkable in Artiodactyl. The number of TRDV1 families in cattle and sheep is 50 and 66 respectively, while TRDV1 is only 13 members in humans. The 29 TRBV genes of Rhinolophus ferrumequinum can be classified into 25 families, and only TRBV6(2 members), TRBV7(2 members) and TRBV12(3 members) are multigene subgroups. According to the statistics of homologous groups of Rhinolophus ferrumequinum (Supplemental Table 5), the amplification of events tends to be random. TRAV24, TRAV25 of horse-headed bat and cattle are obviously polygenic families, but TRAV24, Trav25 of human and dog are monogenic families. Similarly, TRBV6 of horse-headed bat and human are polygenic families, but TRBV6 of dog and pig is monogenic family [10]. TRAV24, TRAV25 of Rhinolophus ferrumequinum and cattle are obviously polygenic groups, but TRAV24, TRAV25 of human and dog are monogenic groups. Similarly, TRBV6 of horse-headed bat and human are polygenic families, but TRBV6 of dog and pig is monogenic family [10]. We counted the CDR region length (Supplemental Table 6) of the homologous TRAV and TRBV genes among different species, and concluded that the CDR region length of the TRBV gene subgroup is conservative among different species, and only the lengths of TRBV1 and TRBV17 are different between human beings and bats. However, the CDR region length of TRAV varies greatly among four species, even though the loss of some statistical information and gene families has affected our statistics. Early researchers have put forward the evolutionary model of IG and TCR genes, which is called: birth-and-death evolution. This evolutionary model explains the emergence of most polygenic families and the fact that some genes become pseudogenes due to extensive mutations, and family amplification occurs at the same time as mutations and deletions occur between families. This is a compensation mechanism to maintain diversity. The internal replication of genome has become a routine event, and the birth of most germline genes comes from the replication of intergenomic fragments, that is to say, the birth of germline genes depends more on the replication of existing genes than on the birth of new subgroups [37-39].

The birth of genes is accompanied by death, because some genes always become pseudogenes due to mutations or frame shifting [39]. 16% of 81 TRAVs, 11% of 18 TRDV genes, 35% of 14 TRGVs and 3% of 29 TRBVs belong to pseudogenes. Correspondingly, the proportion of TRBV pseudogenes in human, dog and pig is 19%, 43% and 31.5% respectively. The TRA/D locus of cattle not only shows high “birth rate”, but also shows high “death rate”. The proportion of TRAV pseudogenes in human, mouse and cattle is 19%, 16.5% and 37.2% respectively [8]. Although there is no diversity degree of cattle, in contrast, 81 TRAV/DV genes of Rhinolophus ferrumequinum show a larger and more stable germline genes pool than 54 TRAV/DV genes of human beings, which is similar to IGVH3 gene family of small brown bat, and there may be more preliminary discoveries of combination diversity [22].

Generally speaking, we want to evaluate the diversity of TCR or BCR receptor repertoire of a species, first from the point of view of the number of germline genes, because this directly affects the diversity of rearrangement, and then consider the addition and mutation of nucleotides. In previous studies, SHM occurred only in B cells of higher vertebrates in order to produce high affinity antibodies, which is very rare in T cells. We counted all species in the IMGT database, including our annotated germline gene pool of Rhinolophus ferrumequinum. Unlike the TRAV gene, most mammals have about 60 TRAJ genes. It is worth noting that sheep and cattle of artiodactyl have a large number of replication events in TRA/D locus and TRAV/dv gene, and the number of TRAJ gene in sheep is 79, but the number of cattle belonging to artiodactyl has not increased significantly. The researchers speculated that the 19 TRAJ complementary genes found in sheep may be the result of replication from TRAJ29 to TRAJ39 due to sequencing errors or amplification [40]. The statistical results of D gene are similar, only cattle and sheep have 9 TRAD genes, and to our surprise, there is only one D gene in Rhinolophus ferrumequinum, which is the only species with only one TRDD gene at present. Another feature of TRG loci is the number of TRGC and the diversity of corresponding protein structures, especially the size of TRGC gene is usually different, and this difference is caused by the different exon number and intron length of coding junction region, which is very obvious even among same species. TRGC2, TRGC3 and TRGC5 of salmon are divided into three exons (EX1, EX2 and EX3), while TRGC1 has two EX2[29]. TRGC1 and TRGC5 of sheep have only one EX2(A), while TRGC3 has two EX2(A and c) and TRGC6 has three EX2(A, b and C) [41]. TRGC5 of cattle has only one EX2, but TRGC3 and TRGC7 have two EX2, while TRGC1, TRGC2, TRGC4 and TRGC6 all have three EX2 [42]. TRGC1 and TRGC2 of dromedary also contain three EX2 [43, 44]. TRGC2, TRGC3 and TRGC4 of dogs have two EX2, while TRGC1, TRGC6, TRGC7 and TRGC8 have only one EX2. All Monoptera and Marsupia lack the second cystamine, which is necessary for the formation of intra-chain disulfide bonds, which is obviously caused by independent mutations [45]. We made a conservative analysis of the RSS sequences of germline genes, but we did not find non-classical RSS before and after the V and J genes without RSS sequences, for example, if the spacer was 12bp±1 or 23bp±1. It would not affect the further analysis, because the effective recombination only occurred between the 12bp RSS and the 23bp RSS [46]. RSS sequence conserved sites of the four loci of Rhinolophus ferrumequinum are all related, whether it is heptamer or nonamer, the first 4 nucleotides of the conserved site of heptamer are CACA, while the conserved site of nonamer is usually the poly-A tract in the sequence. Because the first three nucleotides of heptamer play an important role in the recombination process, the differences of these positions may affect or hinder the recombination of genes, but at present we still don’t know how or how much it will affect the rearrangement if we have atypical RSS before and after germline genes [47].

## 5. Conclusion

Bat is a special kind of mammal, which carries a lot of virulent viruses, but it will not be harmed by these viruses. We still don’t know the mechanism of bat’s anti-virus, not only because of the difficulty in obtaining samples, but also because of the lack of relevant research on bats [17]. In this paper, the four TR loci of Rhinolophus ferrumequinum were annotated completely at the genome level, the structural differences of TR loci between different species and Rhinolophus ferrumequinum were statistically analyzed, and the differences of germline gene composition between different species and Rhinolophus ferrumequinum were discussed. Generally speaking, the four TCR loci of Rhinolophus ferrumequinum are highly conserved compared with other mammals. It should be noted that homologous genes defined by phylogeny may not be able to participate in the rearrangement process, such as homologous genes of TRAV and TRDV, so it is necessary to analyze bat receptor repertoire data through specific primers. On the basis of the annotation of TR loci of Rhinolophus ferrumequinum, our research group has completed the annotation of IG loci of Rhinolophus ferrumequinum (another paper), and has completed the high-throughput sequencing analysis of TCR CDR3 and BCR CDR3 repertoire collected from many types of bats in Xishui area, Guizhou province, China, and compared with the homogeneity and heterogeneity of TCR CDR3 and BCR CDR3 repertoire in physiological conditions, human and mouse central and peripheral, which can provide a basis for studying the acquired immune response mechanism of bats.

## FUNDING

The National Natural Science Foundation of China (31860257); Guizhou Provincial High-level Innovative Talents Project [No. (2018) 5637].

## SUPPLEMENTARY MATERIAL

Statement and acknowledgement

We thank vertebrate genomes project for completing the gene sequencing and sharing the genome sequence. The author of this paper has no conflict of interest and no dispute over his signature.

## Figure legends

Supplementary Figure 1 The IMGT Protein display of the bat complete 81 TRAV genes. Only functional genes ORF and in-frame pseudogenes are shown. The description of the FR-IMGT and CDR-IMGT is according to the IMGT unique numbering for V-REGION

Supplementary Figure 2 The nucleotide and deduced AA sequences of the bat 60 TRAJ genes. The numbering adopted for the gene classification is reported on the left of each gene. The consensus sequences of the J-heptamer and J-nonamer are provided at the figure. The donor splice site for each TRAJ is also shown. The canonical F/W-G-X-G amino acid motifs are shown.

Supplementary Figure 3 Structure of the TRAC、 TRDC and TRBC genes in human (Homo sapiens), rhesus monkey (Macaca mulatta), dog (Canis lupus familiaris), cat (Felis catus), ferret (Mustela putorius furo), rabbit (Oryctolagus cuniculus), sheep (Ovis aries), pig (Sus scrofa) and bat(Rhinolophus ferrumequinum). The numbers correspond to the size of the exons and introns in nucleotides.

**Table. 1.** Statistics analysis of TRA/D loci in human (Homo sapiens), mouse (Mus musculus), cow (Bos taurus), dog (Canis lupus familiaris), cat (Felis catus), rhesus monkeg(Macaca mulatta), sheep (Ovis aries), rabbit (Oryctolagus cuniculus), bat (Rhinolophus ferrumequinum)

| Species | Chromosome Orientation | Size(kb) | Number of TRAV/TRDV genes | Number of TRAJ/TRDJ genes | Number of TRDD genes | Number of TRAC/TRDC genes |
| --- | --- | --- | --- | --- | --- | --- |
| **Homo sapiens** | 14(FWD) | 1000 | 54/8 | 61/4 | 3 | 1/1 |
| **Mus musculus** | 14(FWD) | 1650 | 98/16 | 60/2 | 2 | 1/1 |
| **Bos taurus** | 10(REV) | 3330 | 183/39 | 60/4 | 9 | 1/1 |
| **Canis lupus familiaris** | 8(FWD) | 760 | 58/5 | 59/4 | 2 | 1/1 |
| **Felis catus** | B3(FWD) | 830 | 63/10 | 64/5 | 2 | 1/1 |
| **Macaca mulatta** | 7(FWD) | 806 | 66/5 | 61/4 | 4 | 1/1 |
| **Ovis aries** | 7(REV) | 2880 | 277/70 | 79/4 | 9 | 1/1 |
| **Oryctolagus cuniculus** | 17(FWD) | 900 | 62/4 | 58/3 | 2 | 1/1 |
| **Rhinolophus ferrumequinum** | 6(FWD) | 850 | 84/18 | 60/4 | 1 | 1/1 |

**Table 2.** Statistics analysis of TRG loci in human (Homo sapiens), mouse (Mus musculus), dog(Canis lupus familiaris), cat (Felis catus), rhesus monkey (Macaca mulatta), rabbit (Oryctolagus cuniculus), bat (Rhinolophus ferrumequinum)

| Species | Chromosome<br>Orientation | Size(kb) | Number of<br>TRGV genes | Number of<br>TRGJ genes | Number of TRGC |
| --- | --- | --- | --- | --- | --- |
| **Homo sapiens** | 7(REV) | 160 | 12-15 | 5 | 2 |
| **Mus musculus** | 13(FWD) | 200 | 7 | 4 | 4 |
| **Canis lupus familiaris** | 18(FWD) | 450 | 16 | 16 | 8 |
| **Felis catus** | A2(FWD) | 290 | 12 | 12 | 6 |
| **Macaca mulatta** | 3(FWD) | 145 | 12 | 5 | 2 |
| **Oryctolagus cuniculus** | 10(REV) | 80 | 11 | 2 | 1 |
| **Rhinolophus ferrumequinum** | 20(REV) | 150 | 14 | 6 | 2 |

**Table 3.** Statistics analysis of TRB loci in human (Homo sapiens), mouse (Mus musculus), dog (Canis lupus familiaris), cat (Felis catus), rhesus monkey (Macaca mulatta), rabbit (Oryctolagus cuniculus), bat (Rhinolophus ferrumequinum)

| Species | Chromosome Orientation | Size(kb) | Number of TRBV genes | Number of TRBD genes | Number of TRBJ genes | Number of TRBC |
| --- | --- | --- | --- | --- | --- | --- |
| **Homo sapiens** | 7(FWD) | 620 | 65-68 | 2 | 14 | 2 |
| **Sus crofa** | 18(REV) | 407 | 38 | 3 | 20 | 3 |
| **Canis lupus familiaris** | 16(REV) | 271 | 36 | 2 | 12 | 2 |
| **Felis catus** | A2(FWD) | 302 | 33 | 2 | 12 | 2 |
| **Macaca mulatta** | 3(FWD) | 736 | 77 | 2 | 14 | 2 |
| **Ovis aries** | 4(FWD) | 506 | 94 | 3 | 19 | 3 |
| **Rhinolophus ferrumequinum** | 26(REV) | 203 | 29 | 2 | 15 | 2 |

Data available in IMGT Repertoire(IG and TR)http://imgt.org/IMGTrepertoire/>Locus and genes>Locus description>Locus descriptions>TRA/D, TRG, TRB>Human, Mouse, Cow,
Dog, Cat, Rhesus monkey, Sheep, Rabbit, Pig.

Table S1 Description of the TRA/D genes in the Rhinolophus ferrumequinum chromosome 6 genome assembly (NCBI Reference Sequence NC_046289). The position of all genes and their classification and functionality are reported

Table S2. Description of the TRG genes in the Rhinolophus ferrumequinum chromosome 20 genome assembly (NCBI Reference Sequence NC_046303). The position of all genes and their classification and functionality are reported

Table S3. Description of the TRB genes in the Rhinolophus ferrumequinum chromosome 26 genome assembly (NCBI Reference Sequence NC_046303). The position of all genes and their classification and functionality are reported

Table S4. Description of the TRAV、 TRDV、 TRGV、 TRBV、 TRAJ、 TRDJ and TRBJ ORF and pseudogenes.

Table S5. IMGT Potential germline repertoires of the homologous TRAV subgroups in human (Homo sapiens), dog (Canis lupus familiaris), cow (Bos taurus) and bat (Rhinolophus ferrumequinum). IMGT Potential germline repertoires of the homologous TRBV subgroups in human (Homo sapiens), dog (Canis lupus familiaris), pig (Sus scrofa) and bat (Rhinolophus ferrumequinum),

Table S6. CDR lengths by homologous TRAV subgroups and species in human (Homo sapiens), dog (Canis lupus familiaris), cow (Bos taurus) and bat (Rhinolophus ferrumequinum). CDR lengths by homologous TRBV subgroups and species in human (Homo sapiens), dog (Canis lupus familiaris), pig (Sus scrofa) and bat (Rhinolophus ferrumequinum).

## Notes

### Competing Interest Statement

The authors have declared no competing interest.

