## Supplementary Table for "The TR locus annotation and characteristics of Rhinolophus ferrumequinum"

**Supplementary Table 1.** Description of the TRA/D genes in the Rhinolophus ferrumequinum chromosome 6 genome assembly (NCBI Reference Sequence NC_046289). The position of all genes and their classification and functionality are reported

| Gene identification | Functionality | Position (Complement) |
| --- | --- | --- |
| OG10G2 | ? | 1074…2179 |
| TRAV1 | P | 2456031…2456711 |
| TRAV2-1 | F | 2509279…2509818 |
| TRAV3-1 | F | 2514156…2514626 |
| TRAV2-2 | F | 2520742…2521281 |
| TRAV2-3 | F | 2533230…2533764 |
| TRAV3-2 | F | 2538475…2538942 |
| TRAV4 | F | 2549165…2549970 |
| TRAV6 | F | 2555766…2556329 |
| TRAV13-1 | F | 2565534…2566085 |
| TRAV13-2 | P | 2567730…2568033 |
| TRAV14-1/TRDV1-1 | F | 2571156…2571714 |
| TRAV13-3 | F | 2586064…2586608 |
| TRAV13-4 | F | 2601171…2601712 |
| TRAV14-2/TRDV1-2 | F | 2607284…2607841 |
| TRAV9-1 | F | 2613791…2614288 |
| TRAV13-5 | F | 2623214…2623760 |
| TRAV14-3/TRDV1-3 | F | 2628215…2628773 |
| TRAV9-2 | F | 2634507…2635005 |
| TRAV43 | P | 2640529…2641056 |
| TRAV13-6 | F | 2645433…2645981 |
| TRAV14-4/TRDV1-4 | P | 2652034…2652565 |
| TRAV9-3 | F | 2660373…2660871 |
| TRAV13-7 | F | 2667736…2668277 |
| TRAV14-5/TRDV1-5 | P | 2673428…2673976 |
| TRAV9-4 | F | 2681102…2681598 |
| TRAV13-8 | F | 2689804…2690347 |
| TRAV14-6/TRDV1-6 | F | 2695692…2696250 |
| TRAV11 | ORF | 2701831…2702414 |
| TRAV8-1 | F | 2712087…2712608 |
| TRAV16 | F | 2717620…2718099 |
| TRAV18-1 | F | 2728447…2728911 |
| TRAV19-1/TRDV2 | F | 2730450…2731041 |
| TRAV18-2 | F | 2739487…2739994 |
| TRAV19-2 |  | 2746348…2746606 |
| TRAV20 | P | 2748349…2748677 |
| TRAV21 | F | 2750199…2750776 |
| TRAV8-2 | P | 2753819…2754249 |
| TRAV22-1 | F | 2758182…2758741 |
| TRAV23-1/TRDV3-1 | F | 2767636…2768234 |
| TRAV24-1 | F | 2773550…2774098 |
| TRAV25-1 | F | 2779190…2779820 |
| TRAV8-3 | F | 2786863…2787374 |
| TRAV27-1 | F | 2792990…2793582 |
| TRAV24-2 | F | 2795587…2796135 |
| TRAV25-2 | F | 2802700…2803329 |
| TRAV8-4 | F | 2809777…2810288 |
| TRAV24-3 | F | 2817564…2818106 |
| TRAV25-3 | F | 2824647…2825262 |
| TRAV8-5 | F | 2834937…2835454 |
| TRAV22-2 | P | 2839685…2840243 |
| TRAV23-2/TRDV3-2 | F | 2849279…2849879 |
| TRAV24-4 | F | 2854237…2854785 |
| TRAV25-4 | F | 2859889…2860520 |
| TRAV8-6 | F | 2868335…2868852 |
| TRAV27-2 | F | 2874196…2874789 |
| TRAV24-5 | ORF | 2876405…2876953 |
| TRAV8-7 | P | 2881629…2882094 |
| TRAV22-3 | P | 2884695…2885253 |
| TRAV8-8 | F | 2891305…2891816 |
| TRAV27-3 | F | 2897800…2898392 |
| TRAV24-6 | F | 2900254…2900802 |
| TRAV8-9 | F | 2916784…2917295 |
| TRAV24-7 | P | 2924468…2924992 |
| TRAV25-5 | F | 2931905…2932532 |
| TRAV8-10 | F | 2938623…2939134 |
| TRAV27-4 | F | 2944666…2945259 |
| TRAV29-1/TRDV4-1 | F | 2948007…2948583 |
| TRDV5 | F | 2954415…2955017 |
| TRAV33-1 | F | 2961571…2962173 |
| TRAV8-11 | F | 2971317…2971834 |
| TRAV27-5 | F | 2977378…2977959 |
| TRAV29-2/TRDV4-2 | F | 2980732…2981308 |
| TRAV31 | P | 2984748…2985348 |
| TRAV33-2 | F | 2987593…2988195 |
| TRAV32-1 | F | 2997684…2998499 |
| TRAV34 | P | 3007746…3008383 |
| TRDV6 | F | 3019721…3020314 |
| TRAV32-2 | ORF | 3027843…3028653 |
| TRAV37/TRDV7 | F | 3052797…3053370 |
| TRDV8 | F | 3058179…3058793 |
| TRAV39 | P | 3074862…3075396 |
| TRAV40 | F | 3087363…3087829 |
| TRAV41 | F | 3091326…3091892 |
| TRAV46 | P | 3151979…3152421 |
| TRDV9 | F | 3158158…3158599 |
| TRDV10 | F | 3177016…3177361 |
| TRDD1 | F | 3208329…3208410 |
| TRDJ1 | F | 3209898…3209973 |
| TRDJ2 | F | 3215019…3215093 |
| TRDJ3 | ORF | 3216444…3216526 |
| TRDJ4 | F | 3218774…3218861 |
| TRDC | F | 3222377…3223873 |
| TRDV11 | F | 3228255…3228878 |
| TRAJ1 | ORF | 3234704…3234791 |
| TRAJ2 | F | 3235715…3235799 |
| TRAJ3 | F | 3235963…3236050 |
| TRAJ4 | F | 3237096…3237183 |
| TRAJ5 | F | 3238283…3238373 |
| TRAJ6 | F | 3238929…3239018 |
| TRAJ7 | P | 3241187…3241241 |
| TRAJ8 | F | 3241732…3241827 |
| TRAJ9 | F | 3242377…3242470 |
| TRAJ10 | F | 3245538…3245636 |
| TRAJ11 | F | 3247821…3247896 |
| TRAJ12 | F | 3248687…3248773 |
| TRAJ13 | F | 3249503…3249595 |
| TRAJ14 | F | 3251490…3251577 |
| TRAJ15 | F | 3252059…3252149 |
| TRAJ16 | F | 3252565…3252662 |
| TRAJ17 | F | 3253408…3253498 |
| TRAJ18 | F | 3254474…3254558 |
| TRAJ19 | F | 3255304…3255397 |
| TRAJ20 | F | 3255800…3255889 |
| TRAJ21 | F | 3257506…3257590 |
| TRAJ22 | F | 3259384…3259474 |
| TRAJ23 | F | 3260020…3260109 |
| TRAJ24 | F | 3261551…3261640 |
| TRAJ25 | F | 3262268…3262356 |
| TRAJ26 | F | 3263791…3263879 |
| TRAJ27 | F | 3264856…3264941 |
| TRAJ28 | F | 3265538…3265622 |
| TRAJ29 | F | 3266279…3266372 |
| TRAJ30 | ORF | 3267970…3268027 |
| TRAJ31 | F | 3269807…3269890 |
| TRAJ32 | F | 3270904…3270991 |
| TRAJ33 | F | 3272001…3272094 |
| TRAJ34 | F | 3272651…3272737 |
| TRAJ35 | F | 3274878…3274965 |
| TRAJ36 | F | 3275238…3275324 |
| TRAJ37 | F | 3276091…3276181 |
| TRAJ38 | F | 3276518…3276608 |
| TRAJ39 | ORF | 3278121…3278181 |
| TRAJ40 | F | 3279632…3279714 |
| TRAJ41 | F | 3280368…3280455 |
| TRAJ42 | ORF | 3281360…3281417 |
| TRAJ43 | F | 3281716…3281809 |
| TRAJ44 | F | 3282920…3283010 |
| TRAJ45 | F | 3284305…3284393 |
| TRAJ46 | F | 3285403…3285490 |
| TRAJ47 | ORF | 3286029…3286083 |
| TRAJ48 | F | 3286776…3286866 |
| TRAJ49 | F | 3287660…3287747 |
| TRAJ50 | F | 3288226…3288313 |
| TRAJ51 | F | 3288969…3289061 |
| TRAJ52 | F | 3290824…3290909 |
| TRAJ53 | F | 3291375…3291462 |
| TRAJ54 | F | 3292839…3292294 |
| TRAJ55 | F | 3294244…3294333 |
| TRAJ56 | F | 3295417…3295501 |
| TRAJ57 | F | 3298063…3298153 |
| TRAJ58 | F | 3299045…3299136 |
| TRAJ59 | F | 3299605…3299698 |
| TRAJ60 | F | 3300573…3300662 |
| TRAC1 | F | 3303741…3306544 |
| DAD1 | ? | 3318653…3336546 |

**Supplementary Table 2.** Description of the TRG genes in the Rhinolophus ferrumequinum chromosome 20 genome assembly (NCBI Reference Sequence NC_046303). The position of all genes and their classification and functionality are reported

| Gene identification | Functionality | Position (Complement) |
| --- | --- | --- |
| AMPH | ? | 52424198…52205147 |
| TRGV1-1 | F | 52186520…52186015 |
| TRGV2-1 | P | 52176399…52175881 |
| TRGV3-1 | F | 52162344…52161835 |
| TRGV4-1 | P | 52159300…52158845 |
| TRGV5-1 | ORF | 52155063…52154527 |
| TRGV6-1 | ORF | 52149224…52148706 |
| TRGV7-1 | F | 52147412…52146932 |
| TRGJ1-1 | F | 52140486…52140398 |
| TRGJ2-1 | F | 52138177…52138089 |
| TRGJ3-1 | F | 52134698…52134620 |
| TRGC1 | F | 52131383…52124942 |
| TRGV1-2 | F | 52092696…52092191 |
| TRGV2-2 | P | 52085354…52084836 |
| TRGV3-2 | F | 52074506…52073997 |
| TRGV4-2 | P | 52071457…52071002 |
| TRGV5-2 | ORF | 52067234…52066698 |
| TRGV6-2 | P | 52059010…52058492 |
| TRGV7-2 | F | 52057190…52056702 |
| TRGJ1-2 | F | 52050249…52050161 |
| TRGJ2-2 | F | 52047899…52047811 |
| TRGJ3-2 | F | 52044405…52044328 |
| TRGC2 | F | 52041089…52035058 |
| STARD3NL | ? | 52030211…51976009 |

**Supplementary Table 3.** Description of the TRB genes in the Rhinolophus ferrumequinum chromosome 26 genome assembly (NCBI Reference Sequence NC.046303). The position of all genes and their classification and functionality are reported.

| Gene identification | Functionality | Position (Complement) |
| --- | --- | --- |
| MOXD2 | ? | 8164121…8171343 |
| TRBV1 | F | 8159483…8158814 |
| TRBV3 | F | 8122215…8121717 |
| TRBV4 | F | 8118047…8117546 |
| TRBV5 | F | 8116020…8115522 |
| TRBV6-1 | F | 8088471…8087985 |
| TRBV7-1 | F | 8085469…8084908 |
| TRBV8 | F | 8082269…8081758 |
| TRBV7-2 | F | 8075936…8075431 |
| TRBV13 | F | 8070856…8070332 |
| TRBV6-2 | F | 8066387…8065883 |
| TRBV11 | F | 8062266…8061667 |
| TRBV12-1 | F | 8056213…8055727 |
| TRBV12-2 | F | 8056265…8053779 |
| TRBV12-3 | F | 8052346…8051860 |
| TRBV14 | F | 8048793…8048321 |
| TRBV15 | ORF | 8046948…8046438 |
| TRBV16 | F | 8042654…8042170 |
| TRBV17 | F | 8039889…8039413 |
| TRBV20 | ORF | 8032880…8032160 |
| TRBV21 | F | 8026415…8025921 |
| TRBV22 | F | 8023564…8023053 |
| TRBV23 | F | 8018971…8018477 |
| TRBV24 | F | 8013869…8013368 |
| TRBV25 | F | 8010463…8009929 |
| TRBV26 | F | 8006723…8006205 |
| TRBV27 | F | 8002680…8002167 |
| TRBV28 | F | 7998256…7997750 |
| TRBV29 | F | 7994906…7994271 |
| TRBD1 | F | 7980902…7980824 |
| TRBJ1-1 | F | 7980290…7980216 |
| TRBJ1-2 | F | 7980155…7980081 |
| TRBJ1-3 | F | 7979571…7979495 |
| TRBJ1-4 | F | 7978972…7978895 |
| TRBJ1-5 | F | 7978716…7978640 |
| TRBJ1-6 | F | 7978243…7978164 |
| TRBC1 | F | 7975603…7974159 |
| TRBD2 | F | 7971724…7971643 |
| TRBJ2-1 | F | 7970816…7970740 |
| TRBJ2-2 | F | 7970620…7970543 |
| TRBJ2-3 | P | 7970450…7970403 |
| TRBJ2-4 | F | 7970330…7970255 |
| TRBJ2-5 | F | 7970183…7970109 |
| TRBJ2-6 | F | 7970037…7969960 |
| TRBJ2-7 | F | 7969916…7969842 |
| TRBJ2-8 | F | 7969814…7969735 |
| TRBC2 | F | 7965510…7965176 |
| TRBV30 | F | 7956006…7955267 |
| EPHB6 | ? | 7929436…7911840 |

**Supplementary Table 4**. Description of the TRAV、TRDV、TRGV、TRBV、TRAJ、TRDJ and TRBJ ORF and pseudogenes.

| TRAV | Functionality | Defective Leader | Frameshift | Stop Codon | Defective Splice sites | Defective RSS |
| --- | --- | --- | --- | --- | --- | --- |
| TRAV1 | P |  |  | ● |  |  |
| TRAV8-2 | P |  |  | ● |  | ● |
| TRAV8-7 | ORF |  |  |  |  | ● |
| TRAV11 | ORF | ● |  |  |  |  |
| TRAV13-2 | P | ● | ● | ● | ● | ● |
| TRAV14-4/DV1-4 | P | ● |  | ● |  |  |
| TRAV14-5/DV1-5 | P |  |  | ● |  |  |
| TRAV19-2 | P | ● | ● | ● | ● | ● |
| TRAV20 | P | ● |  | ● |  |  |
| TRAV24-5 | ORF | ● |  |  |  |  |
| TRAV24-7 | P | ● | ● |  |  |  |
| TRAV31 | P | ● |  |  |  | ● |
| TRAV32-2 | ORF | ● |  |  |  |  |
| TRAV34 | P | ● |  | ● |  |  |
| TRAV39 | P |  |  | ● |  |  |
| TRAV43 | P |  |  | ● |  | ● |
| TRAV46 | P |  |  | ● |  | ● |

| TRGV | Functionality | Defective Leader | Frameshift | Stop Codon | Defective Splice sites | Defective RSS |
| --- | --- | --- | --- | --- | --- | --- |
| TRGV2-1 | P | ● |  | ● |  |  |
| TRGV2-2 | P | ● |  | ● |  |  |
| TRGV4-1 | P |  |  | ● |  | ● |
| TRGV4-2 | P |  |  | ● |  | ● |
| TRGV5-1 | ORF | ● |  |  |  |  |
| TRGV5-2 | ORF | ● |  |  |  |  |
| TRGV6-1 | ORF | ● |  |  |  |  |
| TRGV6-2 | P |  |  | ● |  |  |

| J gene | Functionality | Defective motif | Frameshift | Stop Codon | Defective Splice sites | Defective RSS |
| --- | --- | --- | --- | --- | --- | --- |
| TRAJ1 | ORF | ● |  |  |  |  |
| TRAJ7 | P | ● |  |  |  | ● |
| TRAJ30 | ORF |  |  |  |  | ● |
| TRAJ39 | ORF |  |  |  |  | ● |
| TRAJ42 | ORF |  |  |  |  | ● |
| TRAJ47 | ORF |  |  |  |  | ● |
| TRDJ3 | ORF | ● |  |  |  |  |
| TRBJ2-3 | P | ● |  |  |  | ● |

**Supplementary Table 5**. IMGT Potential germline repertoires of the homologous TRAV subgroups in human (Homo sapiens), dog (Canis lupus familiaris), cow (Bos taurus) and bat (Rhinolophus ferrumequinum). IMGT Potential germline repertoires of the homologous TRBV subgroups in human (Homo sapiens), dog (Canis lupus familiaris), pig (Sus scrofa) and bat (Rhinolophus ferrumequinum)

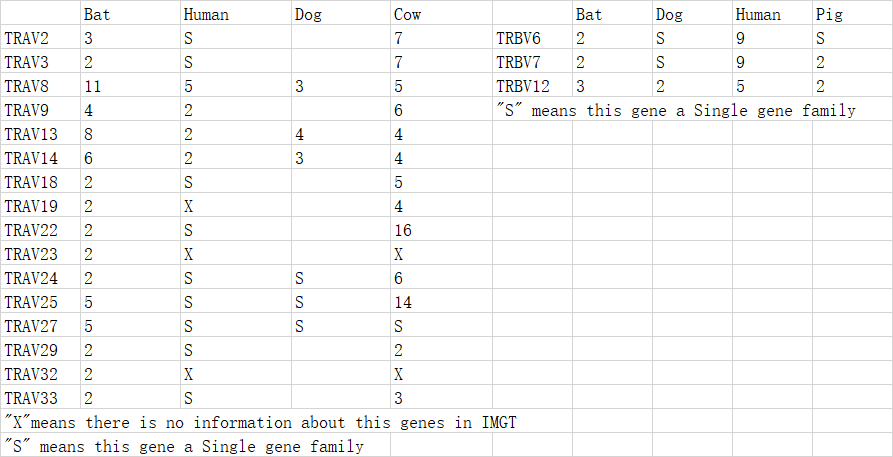

**Supplementary Table 6**. CDR lengths by homologous TRAV subgroups and species in human (Homo sapiens), dog (Canis lupus familiaris), cow (Bos taurus) and bat (Rhinolophus ferrumequinum). CDR lengths by homologous TRBV subgroups and species in human (Homo sapiens), dog (Canis lupus familiaris), pig (Sus scrofa) and bat (Rhinolophus ferrumequinum).

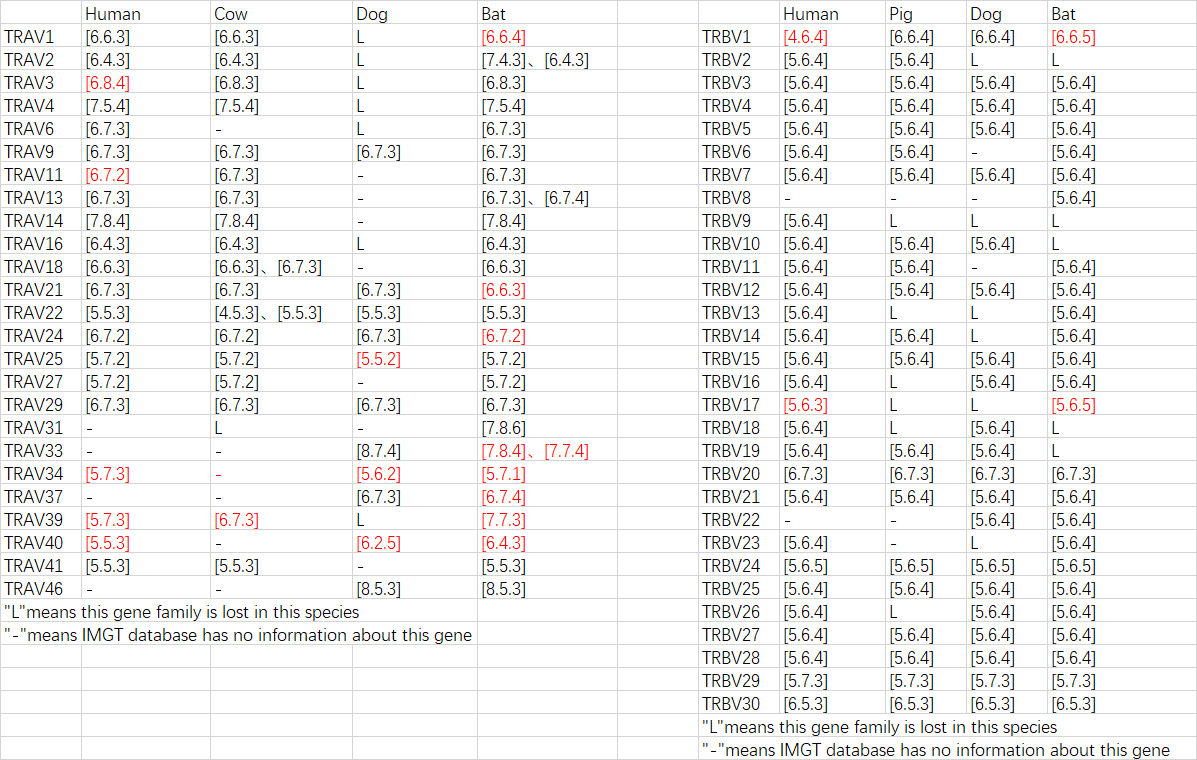
