## Supplementary Figure for "The TR locus annotation and characteristics of Rhinolophus ferrumequinum"

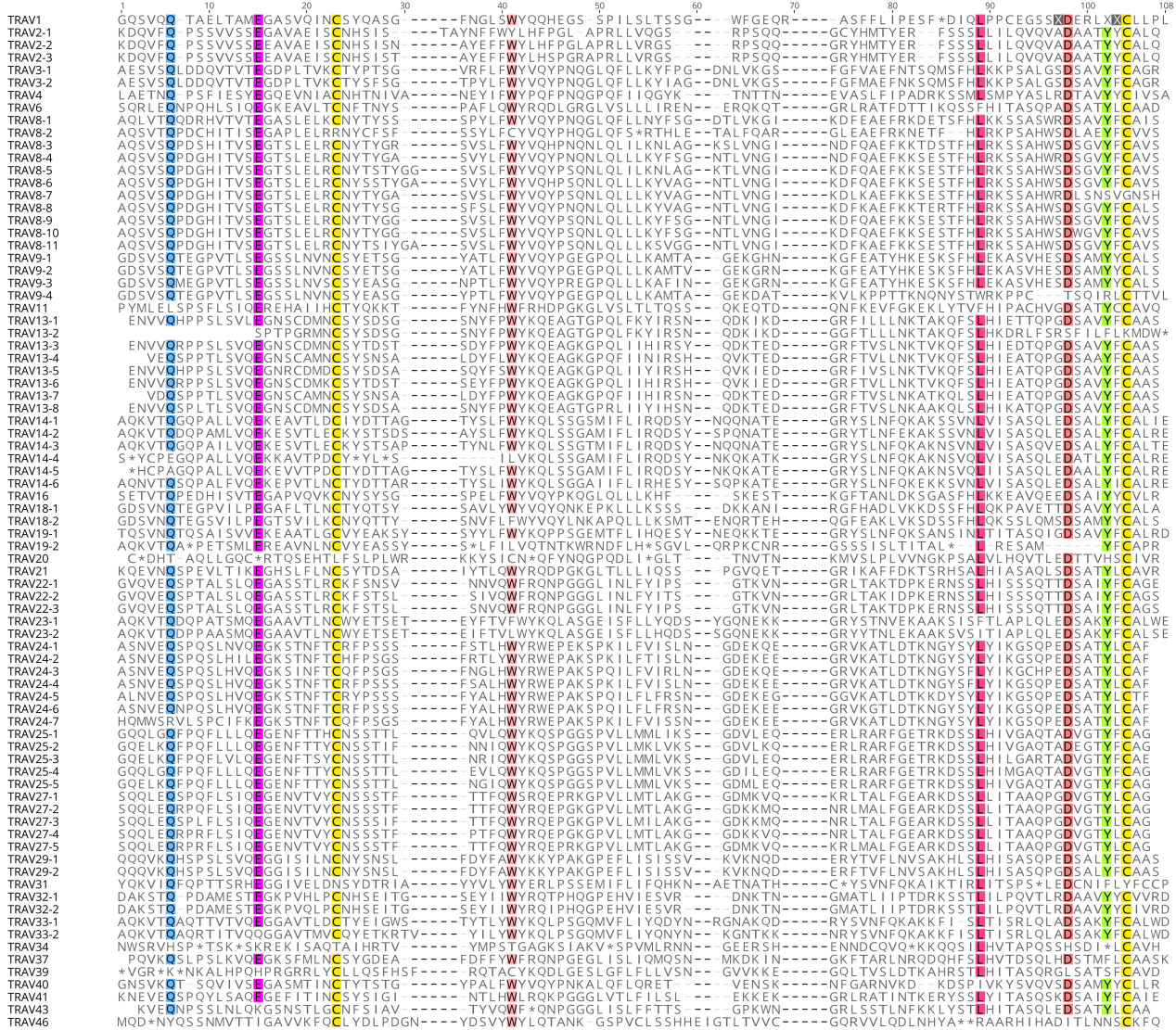

**Supplementary Figure1.** The IMGT Protein display of the bat complete 81 TRAV genes. Only functional genes ORF and in-frame pseudogenes are shown. The description of the FR-IMGT and CDR-IMGT is according to the IMGT unique numbering for V-REGION

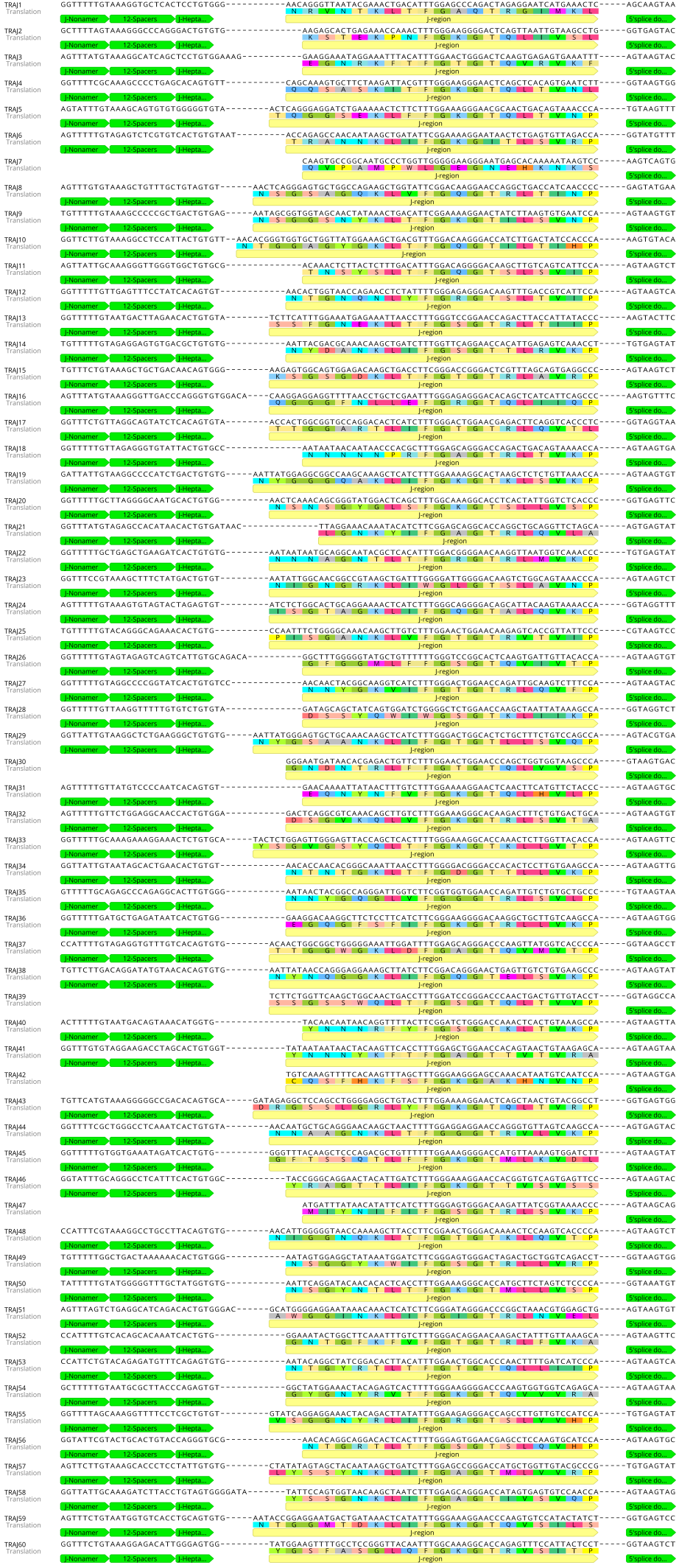

**Supplementary Figure 2.**  The nucleotide and deduced AA sequences of the bat 60 TRAJ genes. The numbering adopted for the gene classification is reported on the left of each gene. The consensus sequences of the J-heptamer and J-nonamer are provided at the figure. The donor splice site for each TRAJ is also shown. The canonical F/W-G-X-G amino acid motifs are shown.

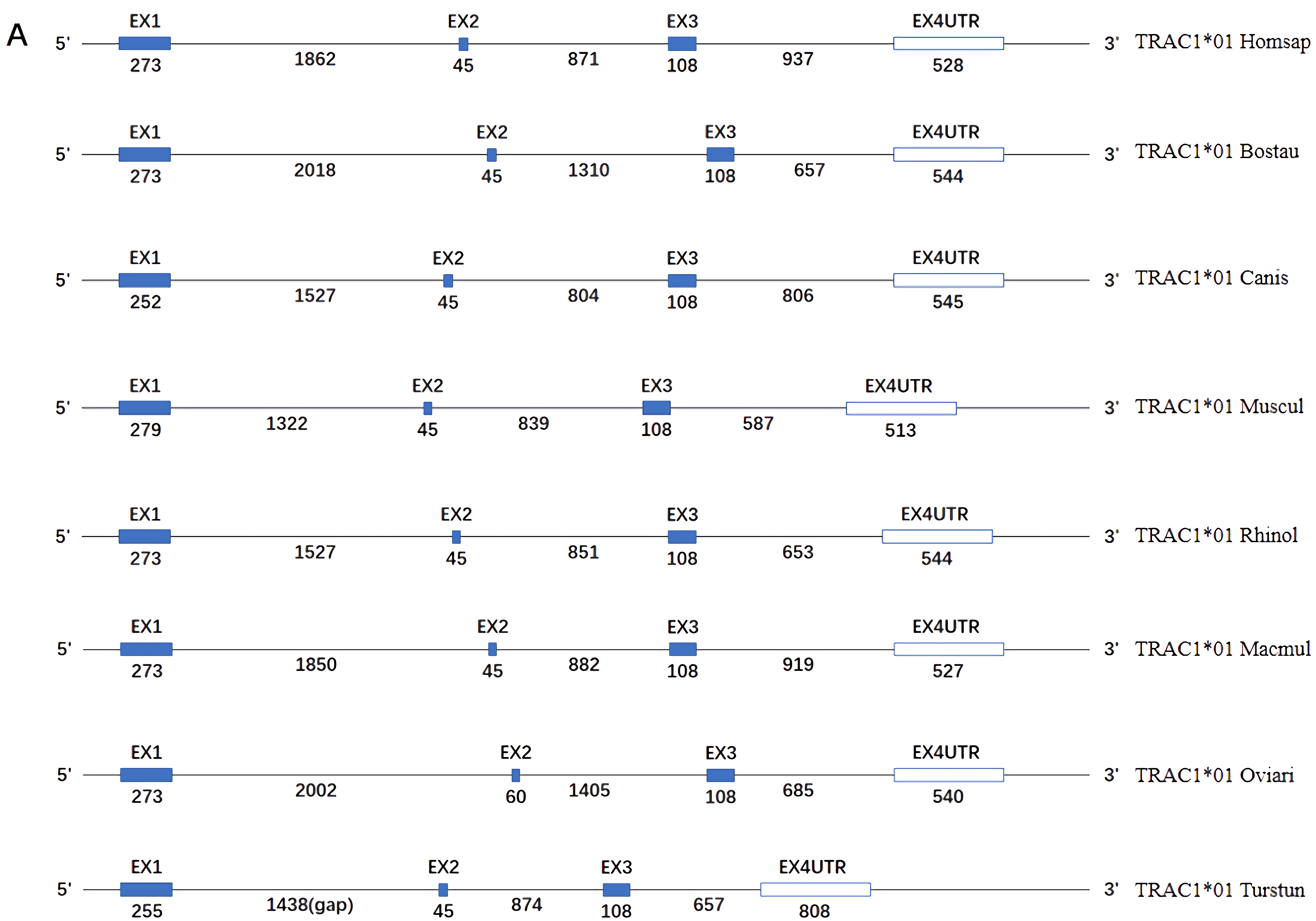

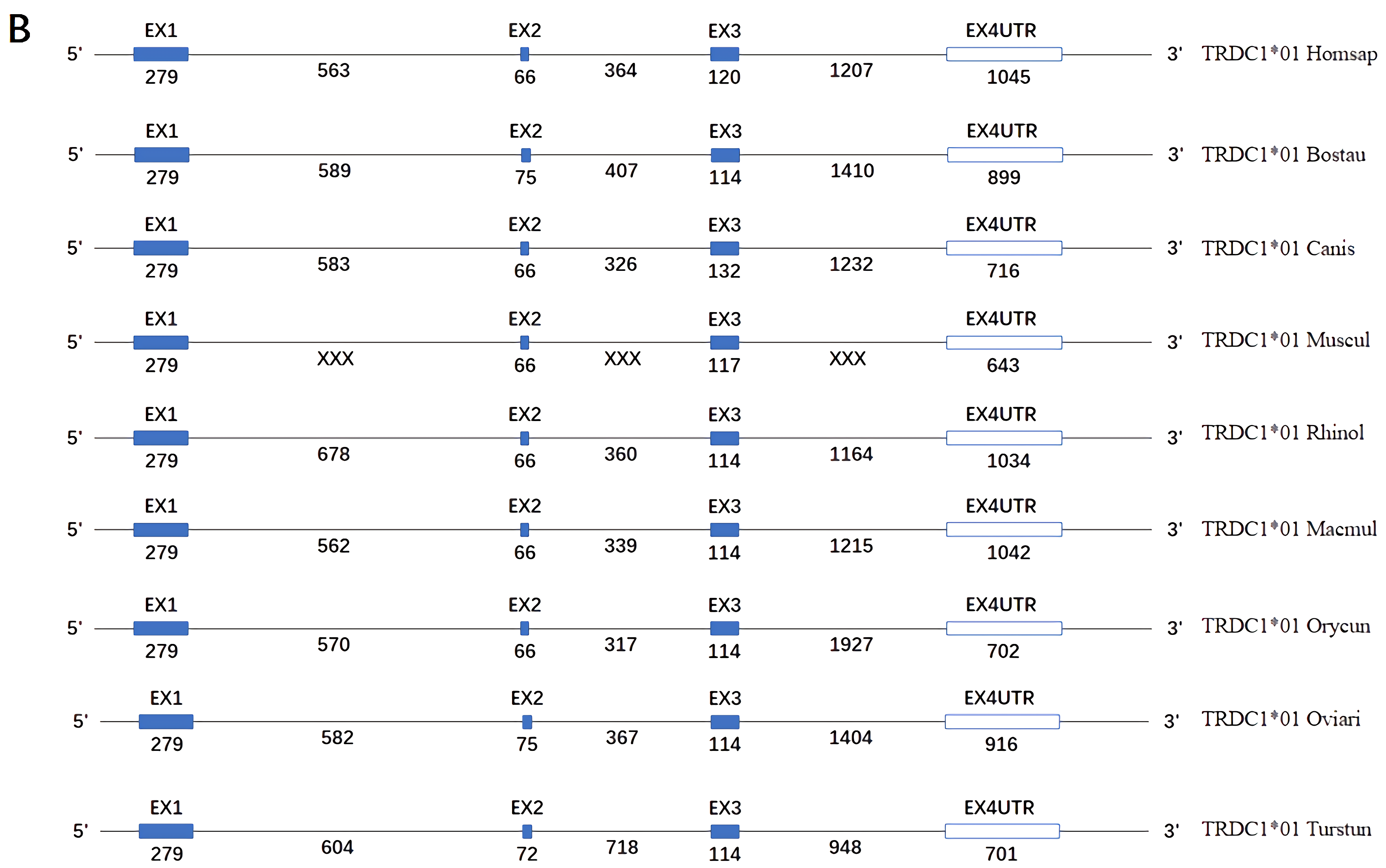

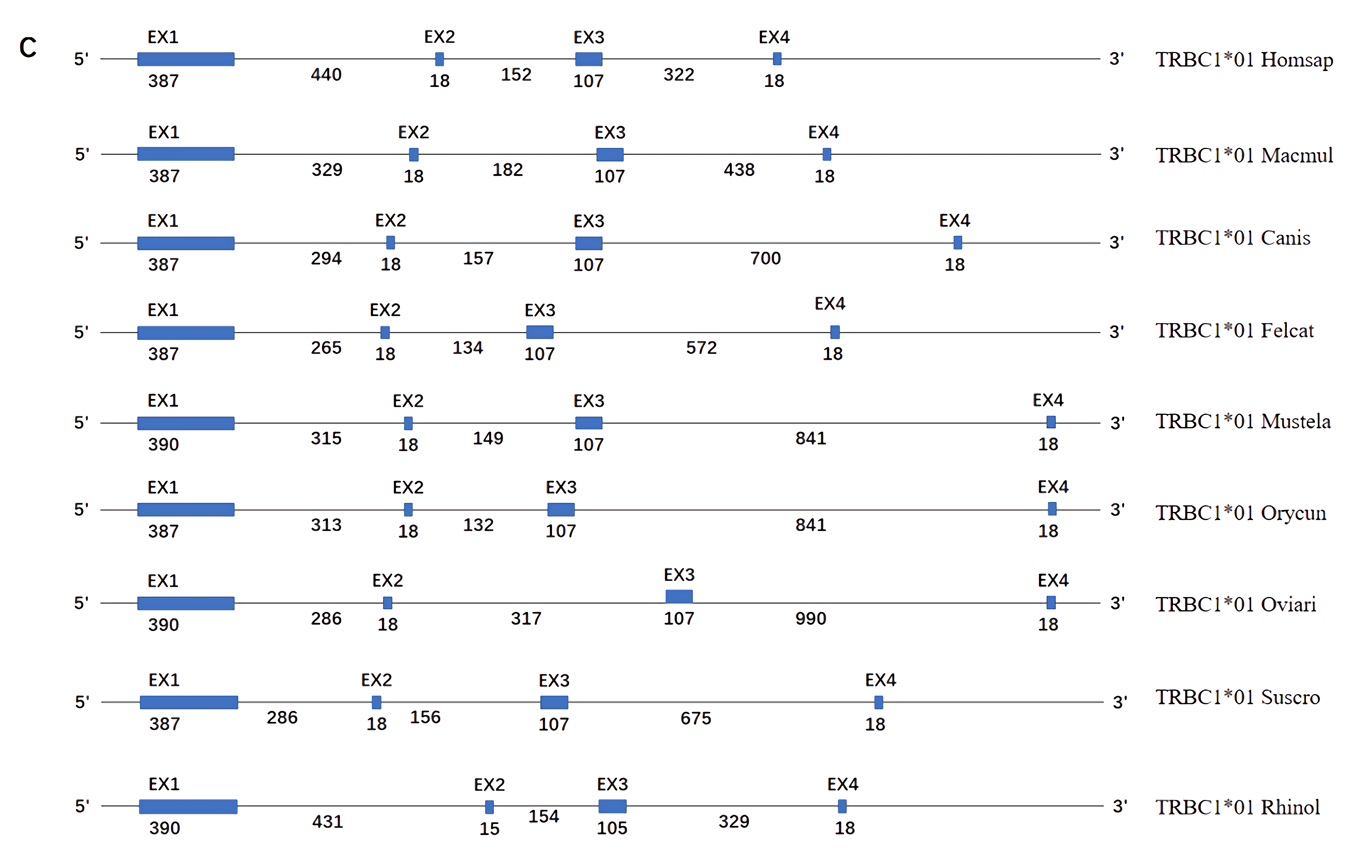

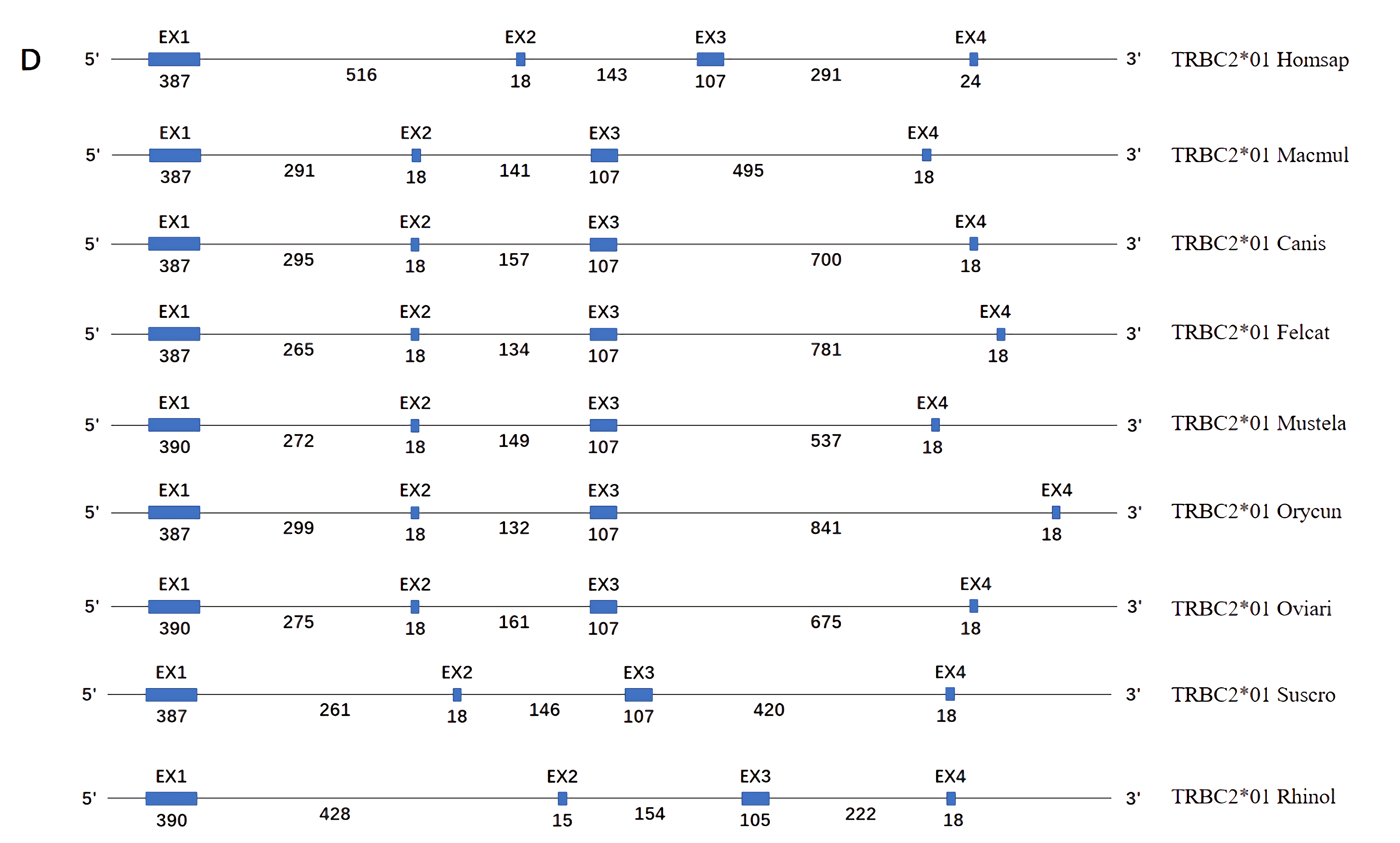

**Supplementary Figure 3.** Structure of the TRAC、TRDC and TRBC genes in human (Homo sapiens), rhesus monkey (Macaca mulatta), dog (Canis lupus familiaris), cat (Felis catus), ferret (Mustela putorius furo), rabbit (Oryctolagus cuniculus), sheep (Ovis aries), pig (Sus scrofa) and bat(Rhinolophus ferrumequinum). The numbers correspond to the size of the exons and introns in nucleotides.
